# Local endoreduplication of the host is a conserved process during Phytomyxea-host interaction

**DOI:** 10.1101/2023.09.21.558765

**Authors:** M Hittorf, A Garvetto, M Magauer, M Kirchmair, W Salvenmoser, P Murúa, S Neuhauser

## Abstract

Endoreduplication is a modified cell cycle in which cells duplicate their DNA without subsequent mitosis. This process is common in plants and can also be found in other organisms like algae and animals. Biotrophic plant pathogens have been shown to induce endoreduplication in their host to gain space and/or nutrients. Phytomyxea (divided into the Plasmodiophorida, the Phagomyxida, and the *Marinomyxa* clade) are obligate biotrophic parasites of plants, diatoms, brown algae, and oomycetes. Here, we tested if phytomyxids induce local endoreduplication in two distant hosts (plants and brown algae). By combining fluorescent in situ hybridisation (FISH) coupled with nuclear area measurements and flow cytometry, we confirmed that endoreduplication is induced by *Plasmodiophora brassicae* (Plasmodiophorida) in infected plants and demonstrate this process in combination with *Maullinia ectocarpii* and *Maullinia braseltonii* (Phagomyxida) in brown algae. We identified molecular signatures of endoreduplication in RNA-seq datasets of *P. brassicae*-infected *Brassica oleraceae* and *M. ectocarpii*-infected *Ectocarpus siliculosus*. Cell cycle switch proteins (CCS52A1 and B in plants and CCS52 in algae) as well as the protein kinase WEE1 (in plants) were identified as genes potentially important for the phytomyxean-induced switch from the mitotic cell cycle to the endocycle. Their expression pattern changed in infected plants and brown algae accordingly. In this study we expand the knowledge on Phytomyxea-host interactions by showing that induced endoreduplication in the host is a conserved feature in phytomyxid infections. The induction of this cellular mechanism by phytomyxid parasites in phylogenetically distant hosts further points at a fundamental importance of endoreduplication in these biotrophic interactions.

## INTRODUCTION

Endoreduplication is a process where the nuclear DNA is multiplied without subsequent cell division (Barlow, 1978; Joubès & Chevalier, 2000), resulting in endopolyploidy where the chromosome number of cells and the cell size increase (Joubès & Chevalier, 2000). Endoreduplication can be found in taxonomically different organisms such as yeasts (Harari et al., 2018), invertebrates (e.g., *Caenorhabditis elegans* (Flemming et al., 2000) and *Drosophila melanogaster* (Smith & Orr-Weaver, 1991)), and in mammals (Gandarillas et al., 2018). Endoreduplication was also described from the green alga *Monostroma angicava* (Horinouchi et al., 2019) and the brown algae *Saccharina latissima*, *Alaria esculenta,* and *Ectocarpus siliculosus* (Bothwell et al., 2010; Garbary & Clarke, 2002). The ultimate reason why many organisms use this alternative cell cycle is still not well understood (De Veylder et al., 2011), but endoreduplication has often been seen as a response to mitigate stress by increasing cell size, gene copy number and levels of gene expression (Paige, 2018; Van de Peer et al., 2021). Endoreduplication is especially widespread in higher plants, where it is important during growth and development (Joubès & Chevalier, 2000; Lee et al., 2009). Cell expansion, cell differentiation and enhanced metabolic activity are processes linked with endoreduplication (Barlow, 1978; Traas et al., 1998) highlighting an essential role during growth and development. In plants, endoreduplication is usually observed when cells shift from cell proliferation and growth to cell differentiation (Joubès & Chevalier, 2000) and is often associated with an increase in cell size and cell expansion (Chevalier et al., 2011; Traas et al., 1998; Wildermuth et al., 2017). In plants endoreduplication plays an important role in endosperm development (Engelen-Eigles et al., 2000; Kowles & Phillips, 1985), but also in processes like overcompensation, i.e., increased numbers of flower, fruit, and seed production under stress conditions (Paige, 2018). Gene expression drastically changes in cells undergoing endoreduplication (Bourdon et al., 2012) as seen in tomato fruits where endoreduplication is hypothesised to increase the metabolic capacity of the plant and to promote growth (Bourdon et al., 2012; Lee et al., 2009; Pirrello et al., 2018). The role of endoreduplication in brown algae however remains unknown, although two reports of endoreduplication exist (Bothwell et al., 2010; Garbary & Clarke, 2002). How the endocycle is induced and regulated in brown algae is also unknown, but some general cell cycle related genes, including cyclins, cyclin dependent kinases, the Wee1 kinase and a cell cycle switch protein (CCS52) homolog, were identified (Bothwell et al., 2010).

The regulation of endoreduplication in plants is better understood than in other organisms. The cell cycle is controlled by oscillating cyclin dependent kinases (CDKs) and their interaction with cyclins (CYC) (de Veylder et al., 2003; Inzé & de Veylder, 2006). Each cell cycle phase (S, M, G1, G2) is regulated by its own specific set of cyclins and CDKs (Komaki & Sugimoto, 2012). For a review of the cell cycle and its detailed regulation in plants see (Berckmans & de Veylder, 2009; Komaki & Sugimoto, 2012; Qi & Zhang, 2020; Shimotohno et al., 2021). For the transition from the regular cell cycle to the endocycle, mitosis specific cyclins and CDKs need to be inactivated (Bhosale et al., 2019). This transition can be regulated/activated through different pathways. In *Arabidopsis thaliana* roots, the endocycle onset is marked by inactivation of the cyclin CYCA2;3 which is controlled through the cell cycle switch protein CCS52A1, an activator of the anaphase-promoting complex (Boudolf et al., 2009). In leaves, CCS52A1 and CCS52A2 seem to control the endocycle through independent pathways (Lammens et al., 2008). The exact role of CCS52B is still unclear, but it is upregulated in nematode infected cells which are in endocycle (De Almeida Engler et al., 2012; Heyman et al., 2017). During leaf development, an alternative way to induce endoreduplication is the downregulation of mitotic CDKs by the SIAMESE-RELATED (SMR) family (cyclin-dependent kinase inhibitors) (Kumar et al., 2015). In tomato fruit, the WEE1 kinase plays a role in endocycle onset, probably through inhibiting CDK activity through phosphorylation (Gonzalez et al., 2004, 2007). For a detailed review about the control and development of endoreduplication in plants see (de Veylder et al., 2011; Lang & Schnittger, 2020).

The obligate biotrophic plant parasitic protist *Plasmodiophora brassicae* induces endoreduplication in infected cells of *A. thaliana* (Olszak et al., 2019). *P. brassicae* belongs to the Phytomyxea (Rhizaria), which are parasites of plants, brown algae, diatoms and oomycetes (Burki et al., 2010; Neuhauser et al., 2011a). The Phytomyxea are divided into the terrestrial Plasmodiophorida, the marine Phagomyxida and the marine Marinomyxa clade (Hittorf et al., 2020; Kolátková et al., 2021). Phytomyxea have a complex life cycle with two phases of infection: the short-lived primary (sporangial) infection and the secondary (sporogenic) infection which often leads to hypertrophy and gall formation in their hosts (Bulman & Neuhauser, 2017; Neuhauser et al., 2011b). During the early phase of secondary infection, the parasite keeps the infected cells in the mitotic cell cycle to promote proliferation (Devos et al., 2006; Malinowski et al., 2019) and carbohydrates are redirected towards the parasite-infected cells (Malinowski et al., 2019). While the secondary plasmodia of *P. brassicae* grow, the host cells change from cell proliferation to cell enlargement and from the mitotic cell cycle to the endocycle (L. Liu et al., 2020; Olszak et al., 2019). This results in local clusters of hypertrophied cells and in *P. brassicae* those are so abundant and large that they lead to the formation of galls in the roots of the host plant (Malinowski et al., 2012). Long lived resting spores are the final stage of the development and those resting spores are eventually released into the soil when the infected roots degrade (Kageyama & Asano, 2009) until the conditions are right and zoospores germinate (Kageyama & Asano, 2009; Wang et al., 2022).

*Maullinia ectocarpii* is an example of a phytomyxean parasite which infects brown algae (Maier et al., 2000). Brown algae are photosynthetic organisms and primary producer in the marine environment which are taxonomically unrelated to angiosperm plants, which makes comparative studies with this parasite compelling from an evolutionary and biological point of view. So far, no spore formation of *M. ectocarpii* has been documented microscopically, although there is persuasive evidence for the presence of the secondary, gall inducing stage on kelp sporophytes (Blake et al., 2017; Mabey et al., 2021). Another clue that *M. ectocarpii* can fulfil a full life cycle is the presence of resting spores in the closely related *Maullinia braseltonii* which commonly infects the bull kelp *Durvillaea* spp. (Murúa et al., 2017). The sporangial phase of the life cycle of *M. ectocarpii* has the advantage that it can be cultured in the lab on suitable brown algal hosts and it is, therefore, available for experimentation (Maier et al., 2000). Brown algae infected with *M. ectocarpii* show hypertrophied infected cells and the host nuclei appear enlarged (Maier et al., 2000), similar to what was found in *P. brassicae* infected plant cells. The biological relevance and the mechanisms which lead to enlarged nuclei and hypertrophy in the host cell are unknown.

Endoreduplication is important during the establishment and maintenance of biotrophic interactions in plants. Many plant biotrophs induce endoreduplication during the colonisation of their host including mutualists, like arbuscular mycorrhizal fungi (AMF) and rhizobia, and parasites, like root knot- and root cyst nematodes or powdery mildew fungi (Carotenuto et al., 2019; Chandran et al., 2010; De Almeida Engler et al., 2012; Fan et al., 2022; Wildermuth et al., 2017). Until now, endocycle induced by symbionts (parasites as well as mutualists) is just recorded in plants and unknown in the interaction between biotrophs and marine brown algae. Endoreduplication and the reprogramming of the host cell cycle have been studied in *A. thaliana* infected with *P. brassicae* (Malinowski et al., 2019; Olszak et al., 2019), while anecdotal evidence in older studies provides information about enlarged host nuclei during infections with the phytomyxids *M. ectocarpii* and *Sorosphaerula veronicae* (Blomfield & Schwartz, 1910; Maier et al., 2000). The study presented here is the first to systematically investigate, quantify and describe the onset and duration of endoreduplication in plant and brown algal hosts during phytomyxean infections.

One aim of this study is to establish whether endoreduplication of infected host cells is a shared, evolutionary conserved mechanism in the class Phytomyxea, used to create space and obtain nutrients and energy from their hosts. To study this, we established a comparative approach, which allowed us to test whether endoreduplication was induced not only by *P. brassicae* in plant hosts but also during the infection of brown algae with *Maullinina spp.* We used a combination of microscopy, ploidy measurements and molecular datasets (RNA-seq) from *P. brassicae* infecting *Brassica spp.* and M. *ectocarpii* infecting *E. siliculosus*; and microscopy observations of *M*. *braseltonii* infecting *D. incurvata* to analyse the induction of endoreduplication. Based on this we provide synergistic evidence supporting the important role of endoreduplication for phytomyxean growth, local energy sink induction and identify new features of their development in plant and brown algae hosts.

## RESULTS

### The shape and size of host nuclei of phytomyxid infected cells are highly variable

The nuclei of uninfected *Brassica rapa subsp. pekinensis* roots were oval to round shaped and differed little in overall size and shape (Figure 1a, Figure 2 a, a’, Supplementary Figure 1 a). The median nuclear area of uninfected plant cells was 19.72 µm^2^ (standard deviation (sd)= 8.5, n=65) with a minimum of 8.76 µm^2^ and a maximum nuclear area of 47.54 µm^2^ (Figure 3a). The median nuclear area of infected *B. rapa* cells was 55.97 µm^2^ (sd=48.9, n=65), the minimum measured was 14.03 µm^2^ and the maximum 278.93 µm^2^ (Figure 3a). The difference between the nuclear area of *P. brassicae* infected cells and uninfected cells was highly significant (based on t-test, p = 0.0000) and the nuclear area of infected cells were 2.8 times bigger based on the median size. While nuclei of normal, uninfected root cells were quite similar in their size and round to oval shaped in their appearance, the nuclei of infected cells varied greatly in their size and had a convex, bulged appearance (Figure 1 a - b, Figure 2 a -d, a’ – d’, Supplementary Figure 3a - b). During the infection of cortical cells of the plant, the plasmodium of *P. brassicae* was gradually growing and occupying more and more space within the host cell. The infected host cells were increasingly hypertrophied, this was accompanied by their nuclei which were gradually getting larger over time (Figure 2 b, b’, c, c’). When the infected cell was completely filled with resting spores of the parasite (Figure 2 d, d’), the host nucleus disappeared.

**Figure 1.**
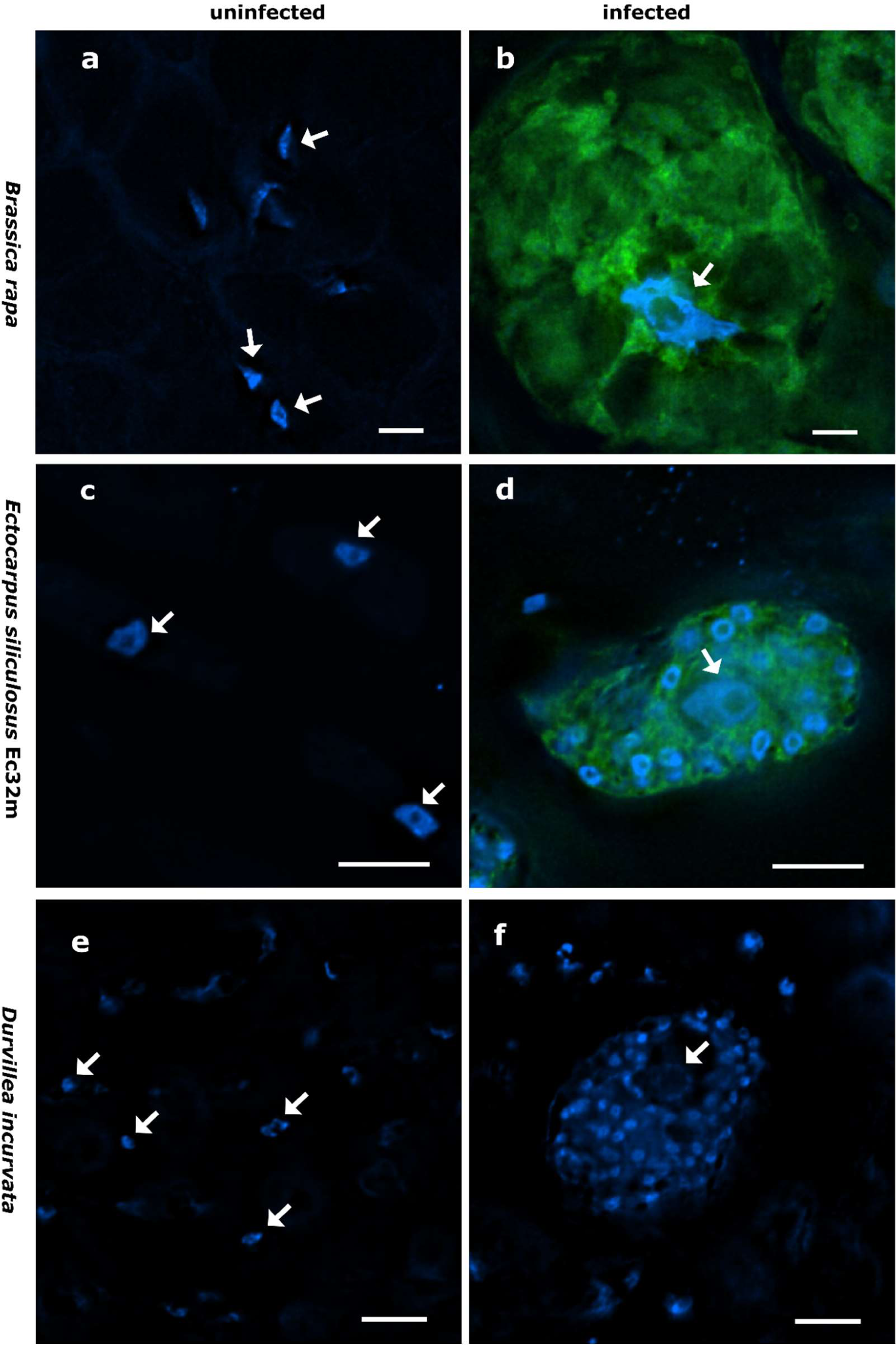
Nucleus size and shape vary between infected and non-infected hosts. Uninfected *Brassica rapa* subsp. *pekinensis* (a), plasmodium of *Plasmodiophora brassicae* in *B. rapa* subsp. *pekinensis* (b), uninfected *Ectocarpus siliculosus* Ec32m (c), multinucleate plasmodium of *Maullinia ectocarpii* in *E. siliculosus* Ec32m (d), uninfected *Durvillea incurvata* (e), and multinucleate plasmodium of *Maullinia braseltonii* in *D. incurvata* (f). Overlay of Hoechst (blue signal, note the smaller nuclei of the phytomyxean plasmodium surrounding the bigger nucleus of the host (arrow)) and FISH (green signal) staining of phytomyxea (a – d); Hoechst staining only (e – f). Arrows point towards host nuclei. Scale bar: 10µm.

**Figure 2.**
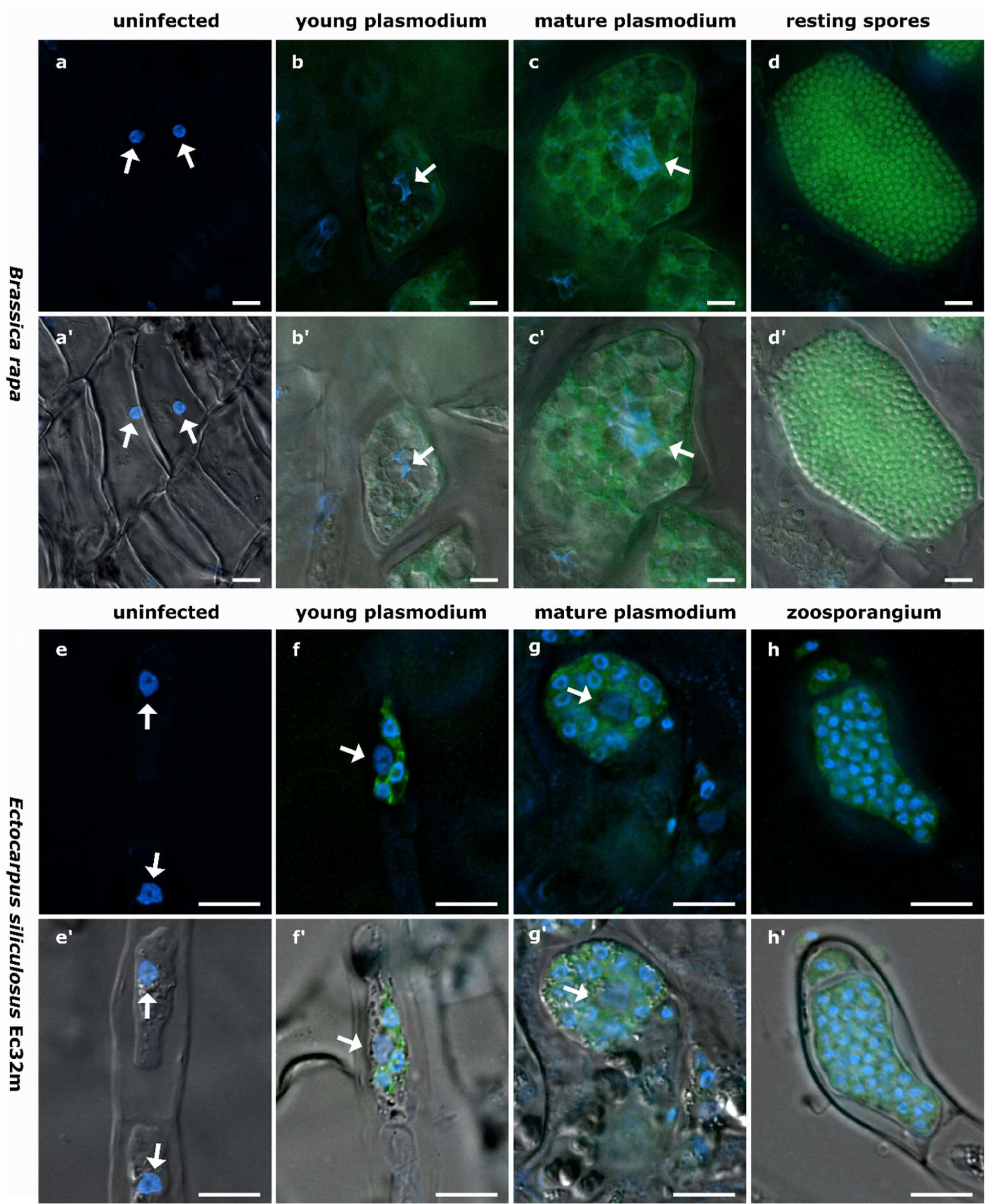
The development of Phytomyxea and the enlargement of host nuclei progress in parallel. *Brassica rapa* infected with *Plasmodiophora brassicae* (a-d, a’ – d’) and *Ectocarpus siliculosus* infected with *Maullinia ectocarpii* (e-h, e’ – h’). Uninfected cells of *B. rapa* (a, a’), young secondary plasmodium of *P. brassicae* (green) in enlarged host cell, host nucleus (arrow) already enlarged (b, b’). Mature secondary plasmodium of *P. brassicae* occupying the now hypertrophied cell and engulfing the enlarged host nucleus (arrow) (c, c’). Uninfected *E. siliculosus* cells (e, e’) in comparison to infected host cells with different infection stages of *M. ectocarpii* (f – h; f’ – h’). Fresh infection of *M. ectocarpii* (green) in *E. siliculosus*, no hypertrophy visible yet (f, f’). Mature plasmodium of *M. ectocarpii*, infected host cell is hypertrophied and the host nucleus is enlarged (arrow) (g, g’). Zoosporangium with zoospores of *M. ectocarpii* occupying the hypertrophied cell of *E. siliculosus*, no host nucleus visible (h, h’). (a – h) Overlay of Hoechst (blue signal, visualized at 365 nm) and FAM (490 nm); (a’-h’) overlay of Hoechst (365 nm), FAM (490 nm) and DIC. *P. brassicae* and *M. ectocarpii* are visualized in green with FISH. Arrows point towards the host nuclei. Scale bars: 10µm.

**Figure 3.**
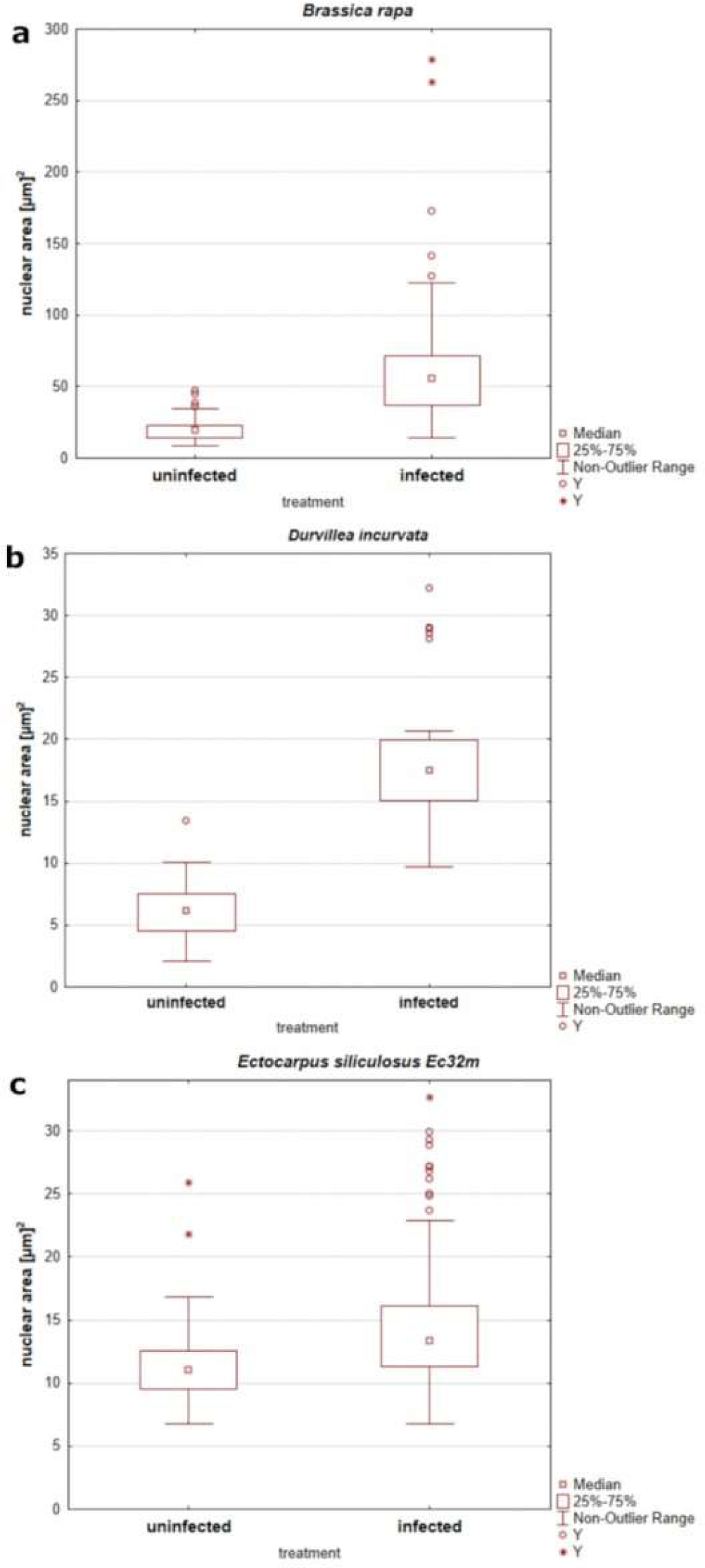
The nuclear area of phytomyxean-infected hosts differs from that of non-infected hosts. (a) Size of the nuclear areas of 65 *P. brassicae* infected *B. rapa* cells and 65 uninfected *B. rapa* cells. (b) Size of the nuclear area of 171 uninfected *E. siliculosus* Ec32m cells compared to the nuclear area of 171 *M. ectocarpii* infected *E. siliculosus* Ec32m cells. (c) Size distribution of 30 uninfected *D. incurvata* cells and 30 *M. braseltonii* infected *D. incurvata* cells.

Nuclei of uninfected *E. siliculosus* (Ec32m) cells had a uniform size and shape (Figure 1 c, Figure 2 e, e’, Supplementary Figure 1 c). The median nuclear area of uninfected *E. siliculosus* cells was 11.03 µm^2^ (sd=2.5, n=171) with a minimum of 6.8 µm^2^ and a maximum of 25.88 µm^2^ (Figure 3 b). The median nuclear area of *M. ectocarpii* infected *E. siliculosus* cells was 13.34 µm^2^ (sd= 4.8, n=171) with a minimum of 6.78 µm^2^ and a maximum of 32.65 µm^2^ (Figure 3 b). Based on the median sizes the nuclei of infected cells were 1.2 times bigger than in uninfected cells and the difference was highly significant (based on t-test, p=0.0000). Uninfected brown algae cells had spherical to ellipsoid nuclei with little variation in size, while nuclei of infected cells showed a high variation in size and a higher variation in shape (Figure 1 c, d, Figure 2 e – h, e’ – h’, Supplementary Figure 1 c, d). The nuclei of infected brown algae were gradually increasing in size along the maturing plasmodium: Cells infected by a young plasmodium still had a normal-sized nucleus (Figure 2 f, f’), whilst cells with a mature plasmodium had enlarged nuclei (Figure 2 g, g’). When the plasmodium differentiated into zoospores and filled the entire host cell, the host nucleus began to disappear (Figure 2 h, h’).

We observed both, plasmodia and resting spores of *Maullinia braseltonii* in the tissue between the cortex and the medulla of infected *Durvillea* blades.The median nuclear area of healthy *Durvillea incurvata* cells was 6.21 µm^2^ (sd= 2.4, n= 30) with a minimum of 2.12 µm^2^ and a maximum of 13.43 µm^2^ (Figure 3 c). The nuclei of *D. incurvata* cells infected with *M. braseltonii* appeared enlarged (Figure 1 f, Supplementary Figure 1 f’). The median nuclear area of infected *Durvillea* cells was 17.51 µm^2^ (sd= 5.8, n=30) with a minimum nuclear area of 9.68 µm^2^ and a maximal nuclear area of 32.15 µm^2^ (Figure 3 c). The difference between infected and uninfected host nuclei was highly significant (based on t-test, p= 0.0000). The nuclei of infected cells were 2.8 times bigger than nuclei of uninfected cells (based on the median of the nuclear area).

TEM images confirmed that the host nuclei of infected *B. rapa* were not apoptotic, but that the increase in size was because of endoreduplication. TEM images showed that the host nucleus of the infected *B. rapa* cell had an intact membrane without holes as would be expected in case of apoptosis. Nucleoli and heterochromatin were present, indicating an active host nucleus (Supplementary Figure 2, Supplementary Figure 3).

Flow cytometry analysis of infected plant and algal material (together with noninfected controls) supported the findings of the nuclear measurements and microscopy. *B. rapa* infected with *P. brassicae* showed ploidy levels of 8C and sometimes a few 16C nuclei were detected (Supplementary Figure 4c, d), whilst roots of control plants had ploidy levels of 4C and rarely 8C (Supplementary Figure 4 a, b). In uninfected *E. siliculosus* samples, only one ploidy level was detected (Supplementary Figure 5 a). In the infected *E. siliculosus* samples, different ploidy levels could be detected, however the interpretation of the peaks must be seen with caution as the peaks are often not well separated and ambiguous (Supplementary Figure 5 b).

### *P. brassicae* infection induces endocycle related transcriptional changes in *Brassica* hosts

We could identify genetic patterns pointing towards the induction of endocycle related processes in *Brassica oleracea var. gongylodes* infected by *P. brassicae* (data from Ciaghi et al., 2019), where transcripts linked to the switch from the mitotic cell cycle to the endocycle (as in Olszak et al., 2019) were generally upregulated. CCS52A1 and CCS52B, activators of the anaphase-promoting complex/cyclosome (APC/C) and involved in the switch from mitotic cycle to the endocycle, were upregulated in infected *B. oleracea var gongylodes* plants confirming the observations in *P. brassicae* infected *A. thaliana* plants from Olszak et al. 2019 (Table 1). CDC20 another activator of the APC/C was upregulated as well and, as a consequence, transcripts of the APC/C were upregulated (Supplementary Table 1). The protein kinase WEE1, which is involved in a different pathway leading to a switch from the mitotic cell cycle to the endocycle, was also upregulated in infected plants (Table 1, Supplementary Table 1).

**Table1.** Changes in activity of cell cycle genes during phytomyxean infection in plant endocycle and brown algae. The genes were detected based on prior literature, particularly when they were linked to biotrophic interactions. Arrows indicate up- or down-regulation of the transcripts in phytomyxea infected plants and algae. Underscore (_) indicates no changes in expression levels. Lack of any symbol indicate absence of the correspondent gene from the dataset. * based on FPKM instead of log2FC; ° (gene names from (Bothwell et al., 2010) and the OrcAE database); °° based on (De Almeida Engler et al., 2012; Gonzalez et al., 2007; Huysman et al., 2015; Inzé & de Veylder, 2006; Joubès & Chevalier, 2000; Lammens et al., 2008; Olszak et al., 2019).

| *E. siliculosus* infected with *M. ectocarpii* |  | *B. oleracea* infected with *P. brassicae* |  | Previously described transcriptional changes °° | function (in plants) | Behaviour of the gene in biotrophic interactions |
| --- | --- | --- | --- | --- | --- | --- |
| gene° | u/d | At homolog /ortholog | u/d |  |  |  |
| CDH1-Ccs52 (Ectsi FZR1) | ↑ | CCS52A1 | ↑* | ↑ | CCS52A1 activator of APC/C; promotes endocycle (Larson-Rabin et al., 2009) | upregulated during <u>nematode</u> infection to promote endoreduplication (De Almeida Engler et al., 2012); upregulated during <u>powdery mildew</u> infection (Chandran & Wildermuth, 2016); homologs of CCS52A are active during <u>rhizobia</u> infection in soybean causing endoreduplication (Fan et al., 2022); Upregulated during <u>*P. brassicae*</u> infection in *A. thaliana* (Olszak et al., 2019) |
|  |  | CCS52A2 |  | ↑ | CCS52A2 activator of APC/C; promotes endocycle (Lammens et al., 2008) | upregulated during <u>nematode</u> infection to promote endoreduplication (De Almeida Engler et al., 2012); homologs of CCS52A are active during <u>rhizobia</u> infection in soybean causing endoreduplication (Fan et al., 2022) |
|  |  | CCS52B | ↑ | ↑ | CCS52B activator of APC/C; promotes endocycle (De Almeida Engler et al., 2012) | upregulated during <u>nematode</u> infection to promote endoreduplication (De Almeida Engler et al., 2012) |
| wee1 (Ectsi Wee1) | ↓ | WEE1 | ↑ | ↑ | WEE1 promotes endocycle in tomato fruit (Gonzalez et al., 2007) | upregulated during <u>*P. brassicae*</u> infection in *A. thaliana* (Olszak et al., 2019) |
| Ectsi CDKA1 | ↑ | CDKA1 | ↑* | ? | control of mitotic cell cycle: G1/S and G2/M (Hemerly et al., 1995; Inzé & de Veylder, 2006) | downregulated during <u>*P. brassicae*</u> infection in *A. thaliana* (Olszak et al., 2019) |
| Ectsi CDKA2/CDKB | ↓ | CDKB1 (;1&2) | ↑_ | ↓ | CDKB1 controls the mitotic cell cycle: G2/M phase specific; negatively regulates the endocycle (Boudolf et al., 2004, 2006; Huysman et al., 2015; Porceddu et al., 2001) | Up and down regulated during <u>*P. brassicae*</u> infection in *A. thaliana* (Olszak et al., 2019) |
|  |  | CDKB2(;1&2) | ↑ | ↓ | control of mitotic cell cycle: G2/M phase specific (transition) (Andersen et al., 2008; Menges et al., 2005) | downregulated during <u>*P. brassicae*</u> infection in *A. thaliana* (Olszak et al., 2019) |
| Ectsi CDKB / 4-like | ↑ |  |  |  |  | - |
| Ectsi E2F | ↑ | E2FA | ↑* | ↑ | E2Fa promotes G1-S transition (Ramirez-Parra et al., 2003; Vandepoele et al., 2005) | upregulated during <u>*P. brassicae*</u> infection in *A. thaliana* (Olszak et al., 2019) |

Transcripts of genes for G1 to S progression (i.e., mainly involved in the replication of the DNA) were upregulated in infected plants compared with uninfected ones (Supplementary Table 1). Transcripts of cyclin D, which are expressed during the G1 and S phase, were upregulated in *B. oleraceae var. gongylodes* (both up and down regulated in infected *A. thaliana* (Olszak et al. 2019)) (Supplementary Table 1). E2Fa, a transcriptional activator of G1/S specific genes, was upregulated in infected plants (Table 1). DPa, its dimerization partner, was also upregulated in infected *B. oleracea* plants (Supplementary Table 1). CDKA important for both the G1/S phase transition and the G2/M phase transition, was upregulated in infected *B. oleracea var. gongylodes* plants (Table 1). Transcripts associated to G2/M specific genes important for the transition to the M-phase did not show a clear pattern of regulation, they were upregulated in infected *B. oleraceae var. gongylodes*; and both up- and downregulated in infected *A. thaliana* (Olszak et al. 2019). G2/M specific CDKB transcripts and cyclins (CYCBs, some CYCAs) were upregulated (Table 1).

### Maullinia ectocarpii infection changes the transcription of genes related to cell-cycle and endocycle in Ectocarpus siliculosus Ec32m

Although the genetic fundamentals of the endocycle in brown algae are not as extensively understood as in plants, a similar function of these genes can be assumed based on homologous genes. Transcripts of Ectsi FZR1 (or CDH1/CCS52), a homolog to the positive endocycle regulator CCS52A in plants, was upregulated in infected *E. siliculosus* (Ec32m) (Table 1). Wee1 (the plant homolog WEE1 is important for endocycle onset in specific tissues in plants) was downregulated in infected algae (Table 1). Transcripts of the Anaphase promoting complex/cyclosome (APC/C), which is activated through CCS52A, were both up and downregulated (Supplementary Table 2).

The expression of genes responsible for regulating the G2-M transition during mitosis was found to be downregulated in infected algae when compared to uninfected hosts. Ectsi CDKA2/CDKB, whose diatom homolog is important for the G2-M transition, was downregulated in infected brown algae (Table 1). Both - A-type cyclins and B-type cyclins - were downregulated in infected algae (Supplementary Table 2).

CDKA1, important for the G1/S transition (i.e., for DNA replication), was upregulated in *M. ectocarpii* infected *E. siliculosus* (Table 1). There was an upregulation of certain transcripts of D-type cyclins, whose plant homologs are involved in the G1/S transition, while others were downregulated. Additionally, E2F, a positive transcriptional regulator of the G1/S transition, was upregulated in infected algae, whilst RBR, its inactivator, was downregulated (Table 1, Supplementary Table 2).

### Phytomyxids may have the ability to control the host cell cycle by using effectors

By analyzing our transcriptomes, we could identify differentially regulated genes of phytomyxids that might act as secreted effectors, influencing the cell cycle of their host (Supplementary Table 3, Supplementary Table 4). Based on TPM values and annotation of the transcripts we identified two potential effectors in *P. brassicae;* a mitotic checkpoint protein (BUB3) and a serine threonine kinase (AURKA). In the *M. ectocarpii* transcriptome a putative Anaphase complex subunit 10 (ANAPC10) and a MOB kinase activator (MOB1) were highly expressed and fulfilled the selection criteria for putative effectors (based on EffectorP).

## DISCUSSION

### Local endoreduplication is a conserved mechanism during phytomyxid-host interaction

During phytomyxid infection, endoreduplication plays a crucial role in creating hypertrophied cells, and consequently allows the parasite to use more space and create a nutrient sink for itself. Both the sporangial phase (*Maullinia ectocarpii*) and the sporogenic phase (*P. brassicae and Maullinia* braseltonii) show endoreduplication, but the effect is stronger during the sporogenic phase. Endoreduplication is an important mechanism involved in plant growth and development (Joubès & Chevalier, 2000; Lee et al., 2009), and changes to the plant endocycle are involved in the successful colonisation of plants by biotrophic pathogens and mutualists (Carotenuto et al., 2019; Chandran et al., 2010; De Almeida Engler et al., 2012; Suzaki et al., 2014). An increase in the size of host nuclei and cells are indicators for active endoreduplication, because DNA in the nucleus is multiplied, but cells do not divide (Carotenuto et al., 2019; Sugimoto-Shirasu & Roberts, 2003). Measurements of the size of the nuclei, analysis of the ploidy of cells and transcriptome data of infected material support our hypothesis that endoreduplication is induced by phytomyxids in their host and that this process is conserved in plants and brown algae. We demonstrate local endoreduplication in plant and stramenopile host cells infected with phytomyxids, hinting at a central role of this altered physiological state of the host cells for the parasite (Figure 4). According to the presented findings, endoreduplication is essential for the growth of all Phytomyxea to induce the energy sink in the host that causes the energy transfer from the host to the parasite, similar to previous findings in *P. brassicae* (Malinowski et al., 2019; Olszak et al., 2019).

**Figure 4.**
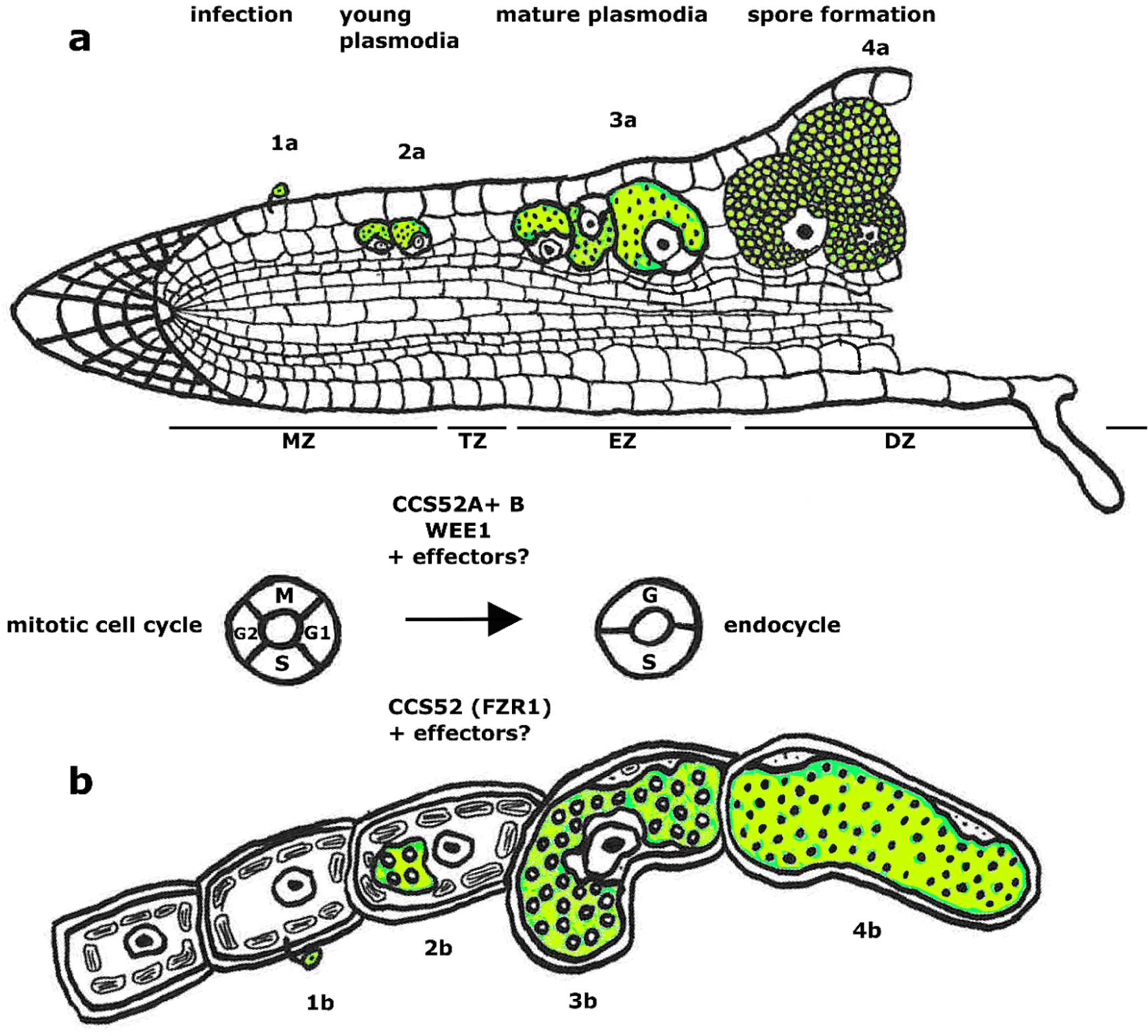
Schematic overview of how phytomyxean infection affects host cell and nucleus size (plant root, a; brown algae, b) Infection process of *P. brassicae* in a plant root from infection with a zoospore (1a), development of multinucleate plasmodia (2a, 3a) to resting spore formation (4a). Below the infection *of M. ectocarpii* in brown algae: 1b infection with a zoospore, development of multinucleate plasmodia (2b, 3b) to formation of a zoosporangium filled with zoospores (4b). The cell cycle machinery of the host switches during the progress of infection (at the plasmodial stage 3a, 3b) from the mitotic cell cycle to the endocycle. The Cell Cycle Switch Proteins (CCS52A1 and CCS52B for plants and CCS52 for algae) play an important role for this switch in both plants and algae. The WEE1 kinase appears to play a significant role in plants.

By examining the changes in infected host cells during the phytomyxid life cycle, it has been observed that the nucleus size of brown algae and plant cells increases in accordance with the growth of phytomyxid plasmodia (Figure 2). The occurrence of endoreduplication is induced when a plasmodium colonizes a host cell and continues until the plasmodium differentiates into resting spores/zoospores (Figure 2), supporting the hypothesis that the induction of local endoreduplication is involved in generating a nutrient sink for the phytomyxid (Malinowski et al., 2019). Plasmodia are the proliferative stage of Phytomyxea, therefore needing a constant energy supply (Garvetto et al., 2023). Cells that undergo endoreduplication show an increase in transcription and metabolic activity, making them an energy sink within the plant (Bourdon et al., 2012; Lang & Schnittger, 2020; Lee et al., 2009). This energy sink is exploited by Phytomyxea until they form resting spores. Microscopic evidence supports the involvement of endoreduplication in feeding the plasmodium as the host nuclei only begin to degrade and disappear during the formation of resting spores of *P. brassicae* and zoospores of *M. ectocarpii* (Figure 2d, h), when metabolic activity in the parasite is likely ceased. Indeed, the nucleus is likely phagocytized by the phytomyxid briefly before the resting spores are differentiated (Garvetto et al., 2023). Furthermore, the similarity in the delayed destruction of the host nucleus in *P. brassicae* sporogenic plasmodia and *M. ectocarpii* sporangial plasmodia is indicative of the importance of endoreduplication in both stages of the life cycle. The source-sink relationship between host and infected cells established in *P. brassicae* (Malinowski et al., 2019) is likely a common, and very basal part of Phytomyxea-host interactions during both intracellular life cycle stages.

### Endoreduplication is a universal, and constant process during the Phytomyxid life cycle

Phytomyxea have two functionally different types of plasmodia: the sporangial (primary) plasmodia are formed during the initial phase of infection and can be found in the main host but also in alternative hosts (Neuhauser et al., 2014; Yang et al., 2022). Sporogenic (secondary) plasmodia, which develop into persistent resting spores and induce large hypertrophied cell clusters, are only found in the main host (Neuhauser et al., 2011a). The data we present indicates that both plasmodia trigger local endoreduplication in their respective host cells, albeit to differing degrees. Sporogenic plasmodia of *P. brassicae* and *M. braseltonii* showed 2.8 times enlarged nuclei in cells where multinucleate parasite plasmodia were present (Figure 1, Figure 3). The sporangial plasmodia of *M. ectocarpii* induced a significant, yet smaller endoreduplication effect on the host than the closely related *M. braseltonii* or *P. brassicae* (1.2 times enlarged nuclei, Figure 1, Figure 3). Because of the limitations in obtaining infected material where both stages are present concurrently, we were unable to measure the effect of both sporangial and sporogenic plasmodia in the same host. Despite this limitation in our study, we hypothesise that different degrees of endoreduplication are linked to biological differences between the two plasmodial stages. The sporangial phase of the phytomyxid life cycle involves relatively short-lived and small plasmodia which colonize the root hairs (in plants) and filamentous thalli/gametophytes (in brown algae) of their host (Kageyama & Asano, 2009; Maier et al., 2000). The sporangial (primary) infection of *P. brassicae* is rapid taking approximately 4-7 days from infection to zoosporangia formation, usually preceding the infection of the cortical cells of the root in the main host (L. Liu et al., 2020). The life cycle of *M. ectocarpii,* which is limited to the sporangial phase in laboratory cultures, takes approximately 14-16 days (personal observations and Maier et al., 2000). In comparison, where known (mainly in *P. brassicae*), the completion of the sporogenic part of the life cycle takes around 20-40 days. The faster transition of sporangial plasmodia from infection to zoospore formation would, therefore, induce a smaller number of endoreduplication cycles, which could explain the findings presented here. The sporogenic part of the phytomyxid life cycle is only found in the main host, where many of the known species of Phytomyxea induce hypertrophy not only at the level of isolated cells, but in large numbers of neighbouring cells resulting in hypertrophied areas and macroscopic galls (Karling, 1968; Kolátková et al., 2021; Murúa et al., 2017; Neuhauser et al., 2010). Infected cells in these galls are filled with large, multinucleate plasmodia with sometimes hundreds of nuclei and it has been estimated that one large clubroot can contain billions of resting spores (Hwang et al., 2013; L. Liu et al., 2020). To produce such large amounts of spores and such large plasmodia, time and energy are needed for the parasite to grow, which is reflected in the longer duration of this part of the life cycle. It remains to be determined whether the even larger host nuclei observed in the sporogenic phase, as compared to the sporangial phase (Figure 3), are directly ascribable to a more specialized interaction between phytomyxid and host; or indirectly caused by the prolonged interaction period between plasmodia and host cells. Regardless, the effect of endoreduplication is more prominent during the secondary phase of the pytomyxid life cycle, especially in gall tissue. It is therefore possible that the two plasmodial stages differs in the way they interact with the host, with the sporangial phase relying on a more transient and less manipulative interaction, than the long-lived specialized sporogenic phase.

The extent of endoreduplication induced by phytomyxids could be influenced by specific limitations to endoreduplication in the host tissue. Indeed, for *P. brassicae* another difference between the sporogenic and sporangial part of the life cycle is the type of host tissue infected, which could also play a role in the host tolerance to increased ploidy in the infected cells. During normal root development at least one round of endoreduplication occurs in the root cortex, which is precisely the tissue usually infected during the sporogenic phase of *P. brassicae* (Bhosale et al., 2019; Pirrello et al., 2018; Zluhan-Martinez et al., 2021). In its sporogenic phase, *M. braseltonii* infects the subcortical area of *Durvillaea spp*. (Murúa et al., 2017), which could be seen as an analogous tissue type in brown algae. The sporangial infection of *P. brassicae* occurs in the root hairs and it is usually only visible during the initial infection (L. Liu et al., 2020), whilst the sporangial phase of *M. ectocarpii* infects growing filamentous thalli or gametophytes of brown algae, i.e., not yet differentiated, young host cells (Maier et al., 2000). Brown algae show differences in their ability to induce endoreduplication as in the filaments of *E. siliculosus* only one round of endoreduplication was observed, while in large kelps with differentiated tissues, more rounds of endoreduplication were observed in the cortex (e.g., *Saccharina latissima*, *Alaria esculenta)* and the medulla (*Saccharina latissima)* (Bothwell et al., 2010; Garbary & Clarke, 2002). Endoreduplication as a conserved feature of phytomyxid-host interactions could have different biological constraints or drivers. Larger plasmodia such as those seen during sporogenic growth could be a result of the increased tolerance of specific tissues for endoreduplication, therefore allowing more energy transfer and longer developmental cycles. It is unclear whether the sporogenic phase is caused by the host ability to tolerate endoreduplication or if it is connected to a specialised manipulation of the host by the parasite, enabling larger plasmodia formation.

### Endoreduplication is caused by modifications in the host gene expression

The microscopic evidence of endoreduplication in infected host cells is supported by transcriptome data, suggesting that Phytomyxea induce endoreduplication through the activation of CCS52 and/or WEE1 kinase (Table 1). In plants, the endocycle can be induced through different pathways including the activation of the APC/C and the consequent inactivation of mitotic cyclins through cell cycle switch proteins (CCS52) in leaves, trichomes and roots (Cebolla et al., 1999; Heyman et al., 2017); the activation of the WEE1 kinase in tomato fruit (Gonzalez et al., 2007); and the inhibition of CDK by SIM/SMR proteins in leaves and trichomes (Kasili et al., 2010; Kumar et al., 2015). Cell cycle switch proteins *CCS52A1* and *CCS52B* were upregulated in infected *B. oleracea var. gongylodes* plants compared to the control plants (Table 1). CCS52A1 was also upregulated during the expansive, late phase of *P. brassicae* infection in *A. thaliana* (Olszak et al., 2019). Homologs of CCS52 were upregulated in *E. siliculosous* infected by *M. ectocarpii,* indicating that CCS52 homologs are induced by Phytomyxea in a conserved manner in plant and stramenopile hosts. Differential expression of CCS52 and homologous genes has been observed in different biotrophic interactions: *CCS52A1*, *CCS52A2* and *CCS52B* are induced in galls during biotrophic nematode infections (De Almeida Engler et al., 2012), the *Medicago* homolog of *CCS52A* is upregulated during arbuscular mycorrhiza symbiosis (Carotenuto et al., 2019), and homologs of *CCS52A* are active in rhizobia-induced nodules in soybean (Fan et al., 2022). This leads to the conclusion that CCS52 is important for the induction of endoreduplication during many biotrophic interactions, especially those that can lead to gall formation in the host.

Transcriptional activation of the WEE1 kinase inhibits CDK activity which subsequently induces endoreduplication (Gonzalez et al., 2007). The regulatory pathway involving the WEE1 kinase was also activated in plant hosts infected with phytomyxids. Transcript levels of WEE1 were increased in *P. brassicae*-infected *B. oleracea var. gongylodes* (Table 1) and *A. thaliana* (Olszak et al., 2019). Although the increase of WEE1 associated transcripts was moderate, it was consistently observed, suggesting that the activation of WEE1 is related to the infection with *P. brassicae*. In *M. ectocarpii-*infected *E. siliculosous*, however, we detected a decrease in expression of *WEE1* homologous transcripts. Endoreduplication induced by *WEE1* is very specific and appears to be restricted to specific tissues or organisms, for example it controls endocycle onset in tomato fruit and maize endosperm, but not in *A. thaliana* leaves (Bhosale et al., 2019; de Veylder et al., 2011). If this activation of WEE1 is tissue and/or species-specific, this could explain the higher ploidy levels observed induced by gall forming Phytomyxea in tissues with the ability to repeat endocycles (de Veylder et al., 2011). However, the role of WEE1 has never been validated in brown algae and might be different; in mammals, for example, WEE1 regulates the cell cycle and is important during DNA damage checkpoints (Elbæk et al., 2020). Ultimately, it can be concluded that the WEE1 kinase-mediated phosphorylation and inactivation of CDKs may have a significant impact on the initiation and continuity of endoreduplication during *P. brassicae* infection, but might have a different role in *M. ectocarpii* infection.

### Multiple factors can lead to the induction of the host endocycle by phytomyxea

How exactly Phytomyxea influence the cell cycle to induce the endocycle is still unknown (Malinowski et al., 2019; Olszak et al., 2019) but our findings allow for a more comprehensive debate. Phytomyxea can induce endoreduplication either actively via effector molecules that target the cell cycle machinery, or passively through the mechanical stress/tension caused by the growing plasmodium (Cao et al., 2017; Toruño et al., 2016). Biotrophs, like arbuscular mycorrhiza fungi, powdery mildews, nematodes and bacteria including rhizobia are known to use effectors to manipulate the host cells (Cornelis, 2006; Goverse & Smant, 2014; Rafiqi et al., 2012; Ratu et al., 2021). The biotroph *Ustilago maydis,* for example, uses an effector to activate mitotic cell division in infected cells and the root-knot nematode *Meloidogyne javanica* likely uses an effector targeting the cell cycle to induce formation of galls (Fitoussi et al., 2022; Redkar et al., 2015). Effectors are key to understand the pathogen-host interaction, therefore potential effectors of *P. brassicae* are well studied (Chen et al., 2019; Muirhead & Pérez-López, 2022; Pérez-López et al., 2020; Rolfe et al., 2016; Schwelm et al., 2015). Our search for effector candidates that interact with the cell cycle revealed a predicted BUB3 (uninhibited by benzimidazole 3) and AURKA (Aurora kinase A) in P. *brassicae* and a predicted APC/C 10 subunit together with a MOB kinase activator (MOB1) in *M. ectocarpii*. BUB3 forms together with BUBR1 and cell division cycle protein 20 (CDC20) the mitotic checkpoint complex (MCC) and also plays a role in phragmoplast-based cytokinesis (Zhang et al., 2018), whilst the AURKA regulates mitosis (Bettencourt-Dias et al., 2004; Karpov et al., 2010). APC/C 10 is a subunit of the anaphase-promoting complex/cyclosome (APC/C) which plays an important role in the regulation of the cell cycle, and together with its activators – the CELL CYCLE SWITCH52 proteins (CCS52A1/A2) – plays an important role in the endocycle in plants (Z. Liu et al., 2019). MOB1 has many roles, also during differentiation and cell progression (Delgado et al., 2020). Those potential effector candidates together with the results of (Pérez-López et al., 2020), who identified a *P. brassicae* cyclin as a putative effector, highlight the potential of effectors in manipulating the host cell cycle. Putative effectors targeting host processes related to the induction of the endocycle are promising targets for future studies to better understand phytomyxid-host interactions.

The second viable hypothesis for the induction of endoreduplication is mechanic stress caused by the growth of the intracellular parasite, indirectly keeping the cells (already undergoing endoreduplication as part of their physiological process of differentiation) “locked” in the endocycle stage until the plasmodium produces spores. Cells in the root cortex are undergoing one or two rounds of endoreduplication during normal root growth (Bhosale et al., 2019). Physical activation of endocycle has been hypothesised for biotrophic nematodes using endocycle competent cells as their feeding sites (de Almeida Engler & Gheysen, 2013). It is also discussed that *P. brassicae* induces wall stress and cell expansion in its host (Badstöber, Ciaghi, et al., 2020), resulting in alterations of the cell cycle (Olszak et al., 2019). Endoreduplication is linked to cell growth and cell wall remodelling (Bhosale et al., 2019; Bhosale & Vissenberg, 2023; Ma et al., 2022), so the growth of the parasite could indirectly induce the host to undergo additional rounds of endocycling, with the resulting energy sink providing nutrients for the Phytomyxea. The causal relationships between endoreduplication, cell growth, and cell wall modifications are not completely understood yet and more studies are needed to identify whether *P. brassicae* induces endoreduplication via mechanical pressure causing cell expansion, or whether cell expansion is a consequence of an effector mediated induction of endoreduplication. The regulation and induction of endoreduplication in brown algae is even less clear, as studies on the topic are scant (Bothwell et al., 2010; Garbary & Clarke, 2002). However, based on the findings discussed here, we hypothesise that similar active and passive processes could be involved in the induction of endoreduplication in all hosts of Phytomyxea.

Local induced endoreduplication is a common feature observed in various plant biotrophic interactions and it is believed that it enhances the proliferation, development, and colonization of biotrophic pathogens. (Wildermuth, 2010; Wildermuth et al., 2017). By locally inducing endoreduplication, an energy sink is generated, most likely providing space for phytomyxid parasites in plant and stramenopile hosts through an increase in cell size and nutrients availability. The capacity of biotrophic parasites to grow and reproduce is constrained by their ability to extract nutrients from the host (Malinowski et al., 2019) and cells undergoing endoreduplication have access to an increased energy pool, as compared to “normal” cells. Indeed, endoreduplication and the related polyploidy is often associated with a more active metabolism (Edgar & Orr-Weaver, 2001; Larkins et al., 2001; Pirrello et al., 2018). Endoreduplication plays a role in cell growth, and ploidy level often correlates with cell size (Breuer et al., 2007; Chevalier et al., 2011; Sugimoto-Shirasu & Roberts, 2003). Increased cell size of the host cells provide space for the growth of the plasmodia, as demonstrated by galls formed in *A. thaliana ccs52a1* mutants with deficient endoreduplication, which were found to be significantly smaller than those of wild type *A. thaliana* (Olszak et al., 2019). Therefore, increased metabolic activity of the host allows for higher energy transfer and combined with larger host cells provides more space and energy to be translated into growth of the phytomyxid parasites. It is yet to be determined whether this interaction occurs as a consequence of indirect processes such as the host’s ability to tolerate endoreduplication and the growth of the plasmodium, which impacts cell growth, or whether it is an active process of communication and negotiation for resources between host and phytomyxid. So far, both possibilities appear plausible based on the available data.

## EXPERIMENTAL PROCEDURES

### Sampling, plant material, algae material and growth conditions

#### Field sampling of infected plant material

*Plasmodiophora brassicae* infected material of *Brassica rapa ssp. pekinensis* was collected in a commercial field in Völs, Tirol at the 17^th^ and 28^th^ of September 2021. Noninfected *B. rapa ssp. pekinensis* samples were harvested in a field in Innsbruck, Tirol at the 30^th^ of September and the 04^th^ of October 2021. Roots were rinsed with tap water and stored at 4 °C until further use as described below.

*Maullinia braseltonii* infected material of *Durvillea incurvata* and healthy *D. incurvata* were sampled at the coast of Estaquilla, Chile at the 19^th^ of May 2022. Samples were cut into 2 cm pieces and fixated with 4% Histofix (phosphate-buffered formaldehyde solution, Carl Roth). Samples were stored at 4°C until further use.

#### Maintenance of *Maullinia ectocarpii* infected *Ectocarpus siliculosus* Ec32m

*Maullinia ectocarpii* (CCAP 1538/1) was grown in *Ectocarpus siliculosus* Ec32m (CCAP 1310/4). Uninfected *E. siliculosus* Ec32m (CCAP 1310/4) was used as a control and grown in the same conditions. Cultures were maintained in artificial sea water with half strength modified Provasoli (West & McBride, 1999) at 15°C with a 12-hour photoperiod, 20 micromol photon m^-2^s^-1^ as described in (Badstöber, Gachon, et al., 2020). Cultures were regularly checked for infections. The cultures were harvested and used in the experiments as described below.

### Preparation of material for microscopy and nuclear measurements

#### Fixation

Plant roots (root galls of *Brassica rapa ssp. pekinensis* infected with *Plasmodiophora brassicae* and roots of uninfected *B. rapa ssp. pekinensis*) were cut with a razor blade and fixated with 4% Histofix (phosphate-buffered formaldehyde solution, Carl Roth) for approx. 1 hour. Afterwards, samples were rehydrated in a series of ethanol washing (10 min 50% EtOH, 2 x 10 min 70 % EtOH, final storage in 100 % EtOH at −20°C). Algal samples (*Maullinia ectocarpii* infected *Ectocarpus siliculosus* Ec32m and uninfected *E. siliculosus* Ec32m) were fixated the same way except for an additional 2.5 min 30% H_2_O_2_ incubation step after fixation in 4 % Histofix to make the cell wall more permeable for the FISH probe.

#### Fluorescence In Situ Hybridization (FISH) and Hoechst staining

To prevent photobleaching, the steps were performed under the protection of red light. Algal samples (infected and healthy *E. siliculosus*) and plant samples (infected and healthy *B. rapa*) were treated in the same way. Fixated samples were incubated for 10 minutes in 35 % hybridization buffer (900 mM NaCl, 20mM Tris HCl, 35% formamide, 0.01% SDS). The hybridization buffer was removed and 100 µl hybridization buffer - probe (**Table 2**; Pl_LSU_2313 for *P. brassicae* and MauJ17 for *M. ectocarpii*) mix (90 µl hybridization buffer and 10 µl probe) was added. The sample was incubated at 46 °C over night. The samples were washed twice with 35% washing buffer (900 mM NaCl, 20 mM Tris HCl, 5mM EDTA, 0.01% SDS) for 20 minutes at 48 °C. For the nuclei staining, the samples were additionally incubated in Hoechst 33342 (Thermo Fisher Scientific, Germany) for 10 minutes and mounted in Vectashield (H-1000, Vector Laboratories). The slide was covered with a coverslip and sealed with nail polish. Slides were stored at – 20 °C in darkness or immediately used.

**Table 2.**
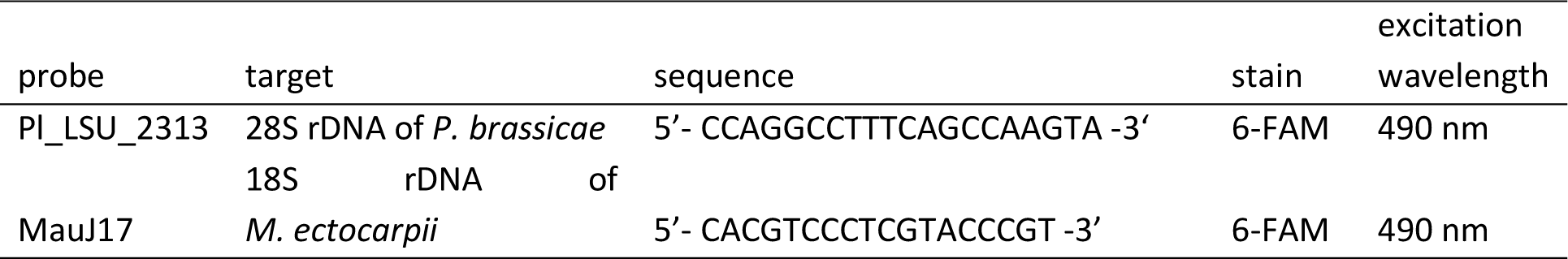
FISH probes used in the experiment, including the target, the sequence, the stain and the excitation wavelength.

Fixated samples of infected and healthy *D. incurvata* were cut into thin sections with a scalpel and stained with Hoechst 33342 (Thermo Fisher Scientific, Germany) for 20 minutes and mounted in Vectashield (H-1000, Vector Laboratories). The slide was covered with a coverslip and sealed with nail polish. Slides were stored at – 20 °C in darkness or immediately used.

### Microscopy

Samples were observed using a Nikon Eclipse Ti2-E (Nikon, Japan) microscope equipped with an Andor Zyla 5.5sCMOS monochrome camera (Andor Technology, United Kingdom) using Nikon CFI Plan-Fluor 40x/0.75 NA and 60x/0.85 NA objectives. The excitation wavelength of 365 nm (for Hoechst 33342) and 490 nm (for the FISH probes) were used. Negative controls without probe but with hybridization buffer and Hoechst 33342 were included. Overlays of the different channels (DIC, channel for Hoechst and channel for the FISH probe) and measurements were done with the NIS Elements software AR 5.21.03 (Nikon, Japan).

### Statistical analysis

Statistica (13.1, StatSoft) was used to analyze the nuclear measurements. A box plot was used to display the differences in the nuclear area of infected vs uninfected host cells. A t – test using Statistica (13.1, StatSoft) was used to test the significance of the differences.

### Transmission electron microscopy (TEM)

Root galls of *Brassica rapa ssp. pekinensis* and roots of noninfected plants were rinsed with tap water. The samples were preselected and screened under the microscope for infections. Selected samples were chemically fixed with 2.5% glutaraldehyde in 0.1M cacodylate buffer containing 10% sucrose at 4 °C for 1h. They were washed with cacodylate buffer and post fixed with 1% osmium tetroxide in 0.05 M cacodylate buffer for 1 h at 4 °C. This was followed by another washing with cacodylate buffer. After dehydration with an increasing acetone series, samples were embedded in Embed 812 resin. A diamond knife (Diatome, Switzerland) and an Ultracut UCT (Leica, Austria) was used to cut cross section of uninfected roots and root galls. The samples were mounted on grids and stained with lead citrate. A Libra 120 energy filter transmission electron microscope (Zeiss, Germany) equipped with a TRS 2×2k high speed camera (Tröndle, Germany) and an ImageSP software (Tröndle, Germany) was used for imaging.

### Flow cytometry

Flow cytometry was performed as described in (Suda et al., 2007) with some modifications explained below.

#### Plants (*Plasmodiophora brassicae* in *Brassica rapa ssp. pekinensis)*

Fresh root galls of *B. rapa ssp. pekinensis* infected with *P. brassicae* were rinsed in tap water. Roots of noninfected *B. rapa spp. pekinensis* were used as a control and treated in the same way. Roots were chopped together with a standard (*Bellis perennis*) in ice cold Otto 1 buffer (0.1 M citric acid, 0.5% Tween 20). The nuclear suspension was filtered through a 42 µm nylon mesh and 1mL Otto2 buffer (0.4 M Na_2_HPO_4_ 12 H_2_O), supplemented with 4’, 6 – diamidino – 2 – phenylindole (DAPI) (Sigma, USA, 4 µg/mL) and 2-mercaptoethanol (2 µl/mL), was added. The relative fluorescence intensity of 3000 particles was measured with a CyFlow space flow cytometer (Sysmex Partec GmbH, Germany) equipped with a UV LED 365 nm. Histograms were analysed using the FloMax software (Partec GmbH, Germany).

#### Algae (*Maullinia ectocarpii* in *Ectocarpus siliculosus*)

*E. siliculosus* Ec32m infected with *M. ectocarpii* was used for flow cytometric measurements. Uninfected *E. siliculosus* Ec32m was used as a control. Fresh algal material was chopped together with the standard *Solanum pseudocapsicum* in modified NIB/2 buffer (NIB: pH 7.5, Sorbitol 125 mM, Potassium Citrate 20 mM, Magnesium Chloride 30 Mm, Hepes 55Mm, EDTA 5mM) supplemented with TritonX-100 (0.1%) and PVP (1%). The nuclear suspension was filtered through a 42 µm nylon mesh and 1mL modified NIB/2 buffer supplemented with TritonX-100 (0.1%), PVP (1%) and DAPI (Sigma, USA, 4 µg/mL) was added. The relative fluorescence intensity of 3000 particles was measured with a CyFlow space flow cytometer (Sysmex Partec GmbH, Germany) equipped with a UV LED 365 nm. Histograms were analysed using the FloMax software (Partec GmbH, Germany).

### Identification of cell cycle related genes in infected hosts

Two publicly available RNA-seq datasets were analyzed to examine the cell cycle-related genes in phytomyxid-infected hosts. The first was from *Brassica oleracea var. gongylodes* infected by *Plasmodiophora brassicae* ((Ciaghi et al., 2019); BioProject: PRJEB26435) and the second from *Ectocarpus siliculosus* Ec32m (strain CCAP 1310/4) infected by *Maullinia ectocarpii* (strain CCAP 1538/1; (Garvetto et al., 2023); BioProject: PRJNA878940). The two transcriptomes (and inferred proteomes) were transformed into blast databases and queried via BLAST (Altschul et al., 1990) for genes (*blastn*) and proteins (*blastp*) homologous to sequences of cell cycle-related genes involved in endoreduplication described by (Olszak et al., 2019) and (Bothwell et al., 2010). Hits were filtered using identity threshold with an identity higher than 50 % for peptides and equal or higher 80% for transcripts and their peptide sequences were blasted against the NCBI database and UniProt database. Resulting gene lists for *B. oleracea* and *E. siliculosus* were compiled and reciprocal keyword searches based on gene models (from *E. siliculosus* to *B. oleracaea* and vice versa) were used to identify additional potential cell cycle-related homologous. Additionally, log2fold change values of infected and uninfected hosts and FPKM values were extracted (Supplementary Table 1, Supplementary Table 2) and analyzed to investigate the behavior of cell cycle-related genes in phytomyxea-infected hosts. The most important genes involved in endoreduplication were summarised in **Table 1** and compared with literature. Potential effectors were identified by filtering the datasets (*M. ectocarpii* and *P. brassicae*) via the cog category D (for cell cycle) together with the EffectorP prediction (only hits from the category effector were kept) Additionally only hits with a peptide identity higher than 50% were kept (Supplementary Table3, Supplementary Table4).

## ACKNOWLEDGEMENTS

This research was funded by the Austrian Science Fund: grant Y0801-B16. Samples from *M. braseltonii* were obtained in the context of the ANID FONDECYT grant 11230059. The authors would like to thank Martin Gachenot for his valuable advice on ploidy measurements in brown algae.

## DATA AVAILABILITY STATEMENT

The data that support the findings of this study are available from the corresponding author upon reasonable request. RNA-seq data are available in the NCBI SRA repository at https://www.ncbi.nlm.nih.gov/sra/ under accession codes PRJEB26435 and PRJNA878940.

## SUPPLEMENTS

**Supplementary Figure 1. Nuclei of host cells infected with phytomyxea.** *B. rapa* subsp. *pekinensis* (a) and infected with *P. brassicae* (b), *E. siliculosus* Ec32m (c) infected with *M. ectocarpii (d), D. antarctica* (e) infected with and *M. braseltonii* (e). Overlay of DIC image, Hoechst staining and FISH (a - d); and overlay of DIC image and Hoechst (e, f). Arrows point toward host nuclei. Scale bar: 10µm.

**Supplementary Figure 2. Transmission electron microscopy (TEM) of *B. rapa subsp. Pekinensis infected with Plasmodiophora brassicae***. Infected host cell with an intact enlarged host nucleus (arrowhead) surrounded by the plasmodium of *P. brassicae* (a, a’). Uninfected root cell of *B. rapa subsp. pekinensis* with smaller, well defined nuclei (b, b’). Scale bar 2 µm. Annotations: HN Host nucleus, Hn Host nucleolus, PN parasite nucleus. Original images (a, b) and images with added annotations to clarify the important features (a’, b’).

**Supplementary Figure 3. Detailed TEM image of *B. rapa subsp. Pekinensis infected with P. brassicae.*** Infected host cell with an intact enlarged host nucleus surrounded by the plasmodium of *P. brassicae* (a, a’). Uninfected root cell of *B. rapa subsp. pekinensis* shows a “normal” sized host nucleus (b, b’). Scale bar 2 µm. HN Host nucleus, Hn Host nucleolus, Ec Euchromatin, Hc Heterochromatin, HMt Host mitochondrion, CW cell wall

**Supplementary** Figure 4. Histogram of relative DNA content from flow cytometry data of (a, b) uninfected control *Brassica rapa* roots and (c, d) *Plasmodiophora brassicae* infected *Brassica rapa* roots (infected, second row). S..standard (*Belli perennis*), B.r… *Brassica rapa*.

**Supplementary Figure 5.** Histogram of relative DNA content from flow cytometry data of (a) uninfected *Ectocarpus siliculosus* Ec32m and (b) *Maullinia ectocarpii* infected *Ectocarpus siliculosus* Ec32m cells. S..standard (*Solanum pseudocapsicum*), Ec32m…*Ectocarpus siliculosus* Ec32m, Me *Maullinia ectocarpii*.

**Supplementary Table 1.** Differentially expressed cell cycle genes in *P. brassicae* infected *B. oleracea.* Data from white galls, which are galls that still show considerable growth of *P. brassicae* where compared to noninfected *Brassica* roots. Upregulated genes (↑), downregulated genes (↓). Log2fold changes highlighted using a colour gradient (upregulated in green to downregulated in red). Genes with a key role in endoreduplication are highlighted in blue.

**Supplementary Table 2.** *E. siliculosus* (Ec32m) cell cycle genes (according to Bothwell et al. 2010) differentially expressed when infected with *M. ectocarpii*. Upregulated genes (↑), downregulated genes (↓). Log2fold changes highlighted using a colour gradient (upregulated in green to downregulated in red). Genes important for the switch from the mitotic cell cycle to the endocycle are highlighted in blue.

**Supplementary Table 3.** Putative *M. ectocarpii* effectors with predicted functions involved in the cell cycle (cog cat = D).

**Supplementary Table 4.** Putative effectors of *P. brassicae* which are predicted to be involved in the cell cycle.

