## Supplementary Figure for "Local endoreduplication of the host is a conserved process during Phytomyxea-host interaction"

**Supplementary Figures 1- 5**

**Supplementary Tables 1-4**

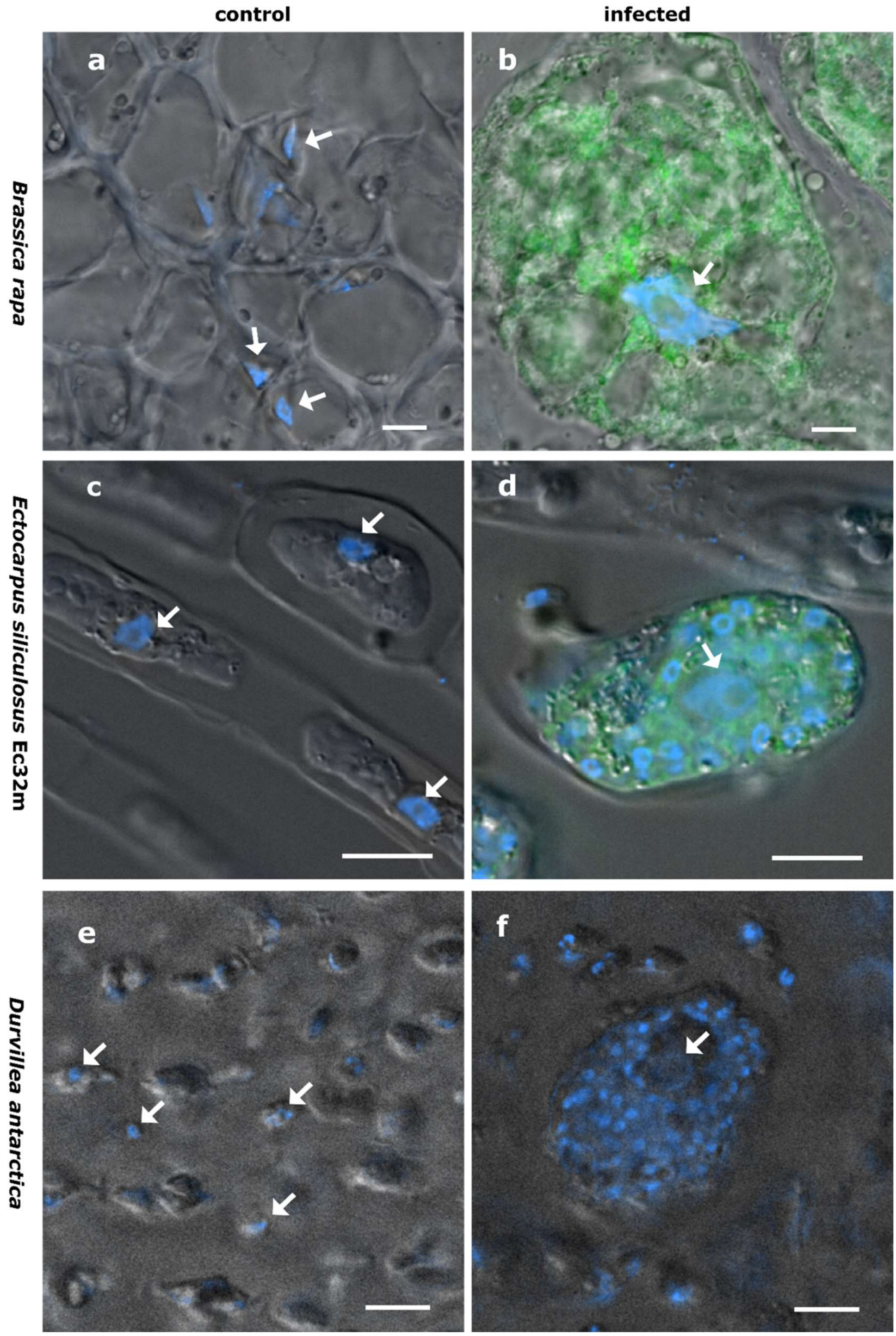


**Supplementary Figure 1. Nucleus size and shape vary between infected and non-infected hosts.** Uninfected *Brassica rapa subsp. pekinensis* (a), plasmodium of *Plasmodiophora brassicae* in *B. rapa subsp. pekinensis* (b), uninfected *Ectocarpus siliculosus* Ec32m (c), multinucleate plasmodium of *Maullinia ectocarpii* in *E. siliculosus* Ec32m (d), uninfected *Durvillea incurvata* (e), and multinucleate plasmodium of *Maullinia braseltonii* in *D. incurvata* (f). Overlay of DIC image, Hoechst and FISH (a, b, b, d); overlay of DIC image and Hoechst (e, f). Scale bar: 10µm.

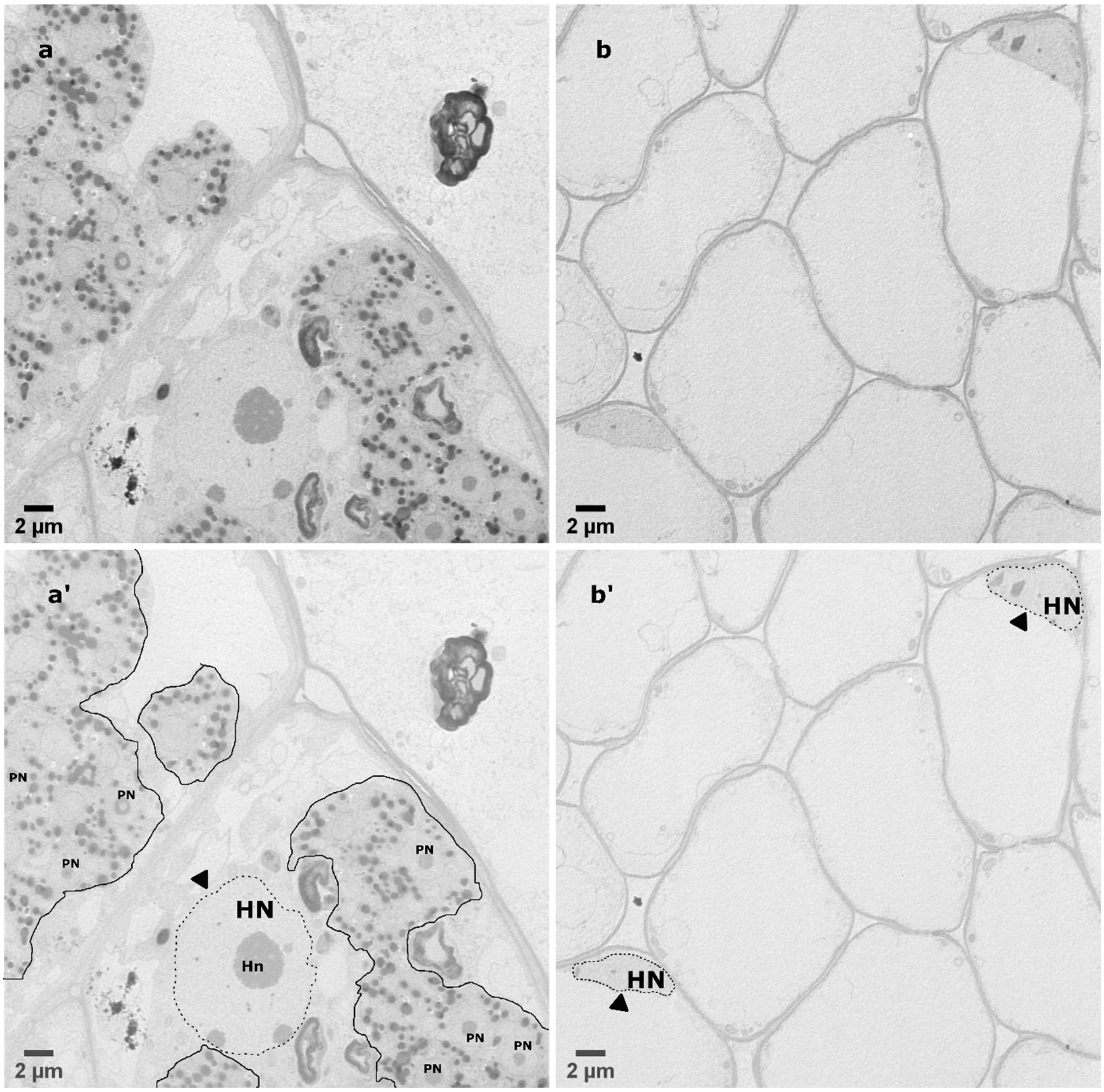


**Supplementary Figure 2. Transmission electron microscopy (TEM) of *Plasmodiophora brassicae* infected *Brassica rapa subsp. pekinensis* and uninfected *B. rapa subsp. pekinensis* root cells.** Infected host cell with an intact enlarged host nucleus (arrowhead) surrounded by the plasmodium of *P. brassicae* (a, a’). Uninfected root cell of *B. rapa subsp. pekinensis* shows a “normal” sized host nucleus (b, b’). Scale bar 2 µm. HN Host nucleus, Hn Host nucleolus, PN parasite nucleus.

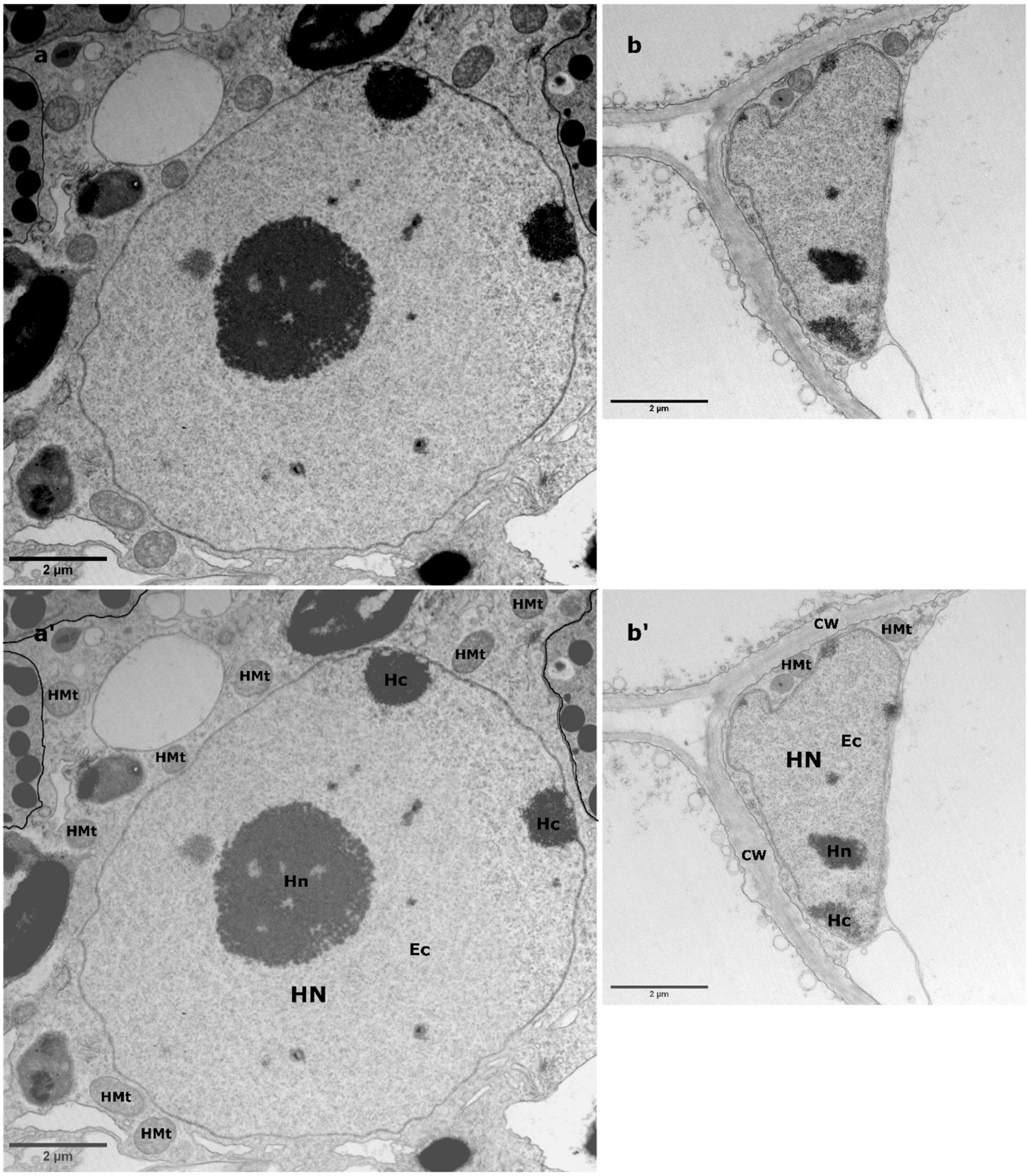


**Supplementary Figure 3. Transmission electron microscopy (TEM) of *Plasmodiophora brassicae* infected *Brassica rapa subsp. pekinensis* and uninfected *B. rapa subsp. pekinensis* root cells.** Infected host cell with an intact enlarged host nucleus (arrowhead) surrounded by the plasmodium of *P. brassicae* (a, a'). Uninfected root cell of *B. rapa subsp. pekinensis* shows a "normal" sized host nucleus (b, b'). Scale bar 2 µm. HN Host nucleus, Hn Host nucleolus, Ec Euchromatin, Hc Heterochromatin, HMt Host mitochondrion, CW cell wall

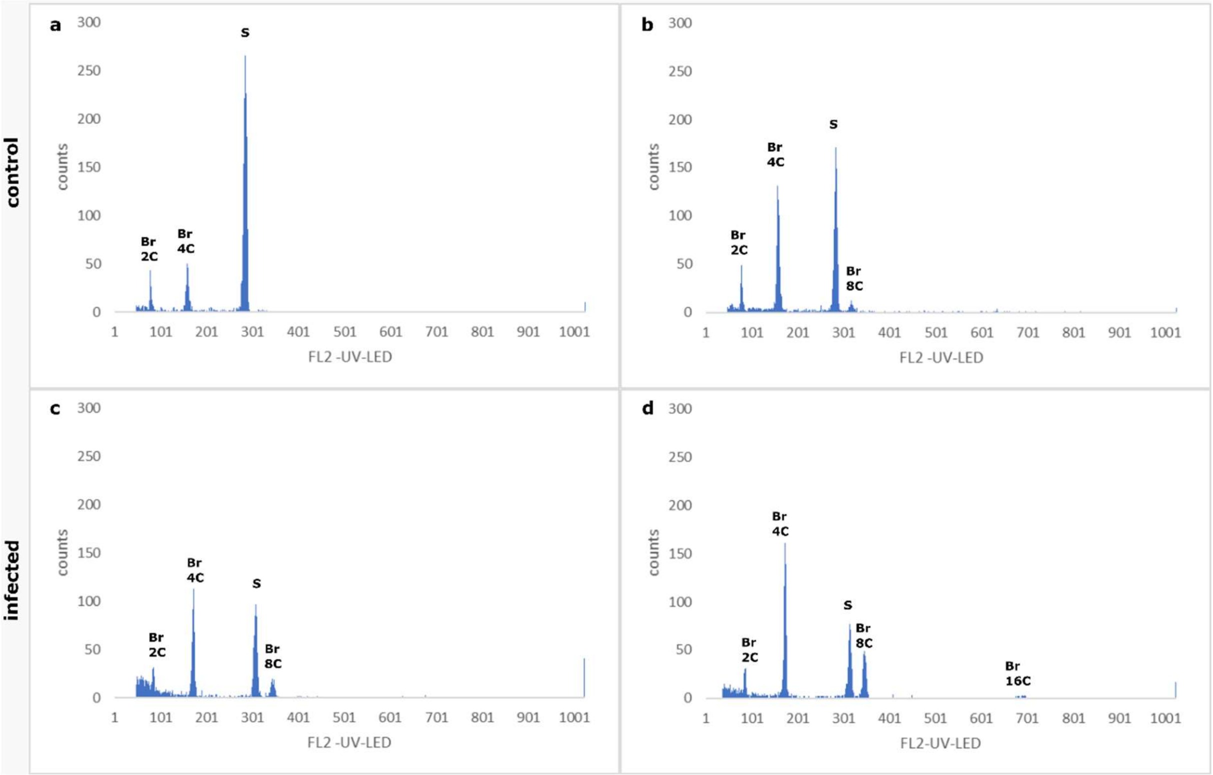


**Supplementary Figure 4.** Histogram of relative DNA content from flow cytometry data of (a, b) uninfected control *Brassica rapa* roots and (c, d) *Plasmodiophora brassicae* infected *Brassica rapa* roots (infected, second row). S..standard (*Belli perennis*), B.r… *Brassica rapa*.

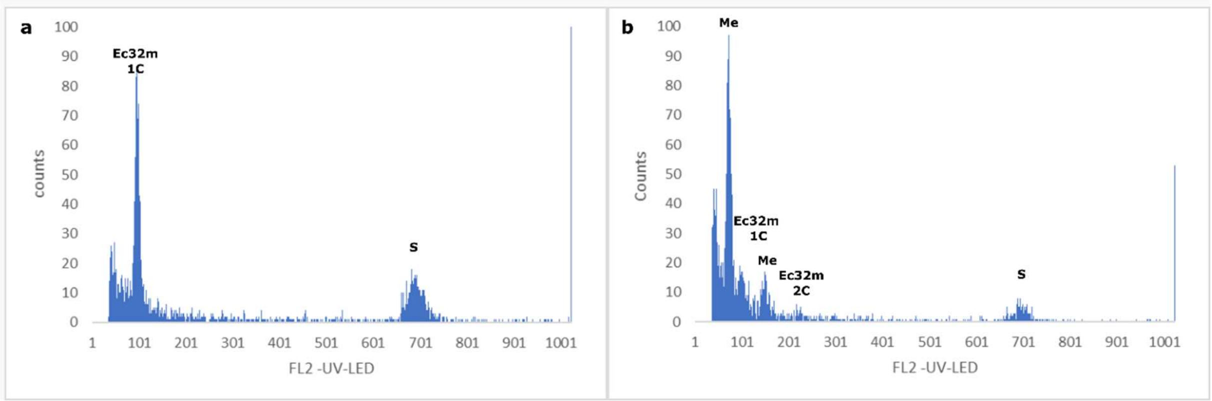


**Supplementary Figure 5.** Histogram of relative DNA content from flow cytometry data of (a) uninfected *Ectocarpus siliculosus* Ec32m and (b) *Maullinia ectocarpii* infected *Ectocarpus siliculosus* Ec32m cells. S..standard (*Solanum pseudocapsicum*), Ec32m…*Ectocarpus siliculosus* Ec32m, Me *Maullinia ectocarpii*.

**Supplementary Table 1.** Cell cycle genes differentially expressed in *Plasmodiophora brassicae* infected *Brassica oleracea* in comparison to noninfected *Brassica oleracea*. White galls where used for the comparison with the noninfected *Brassica* roots. Upregulated genes are highlighted with an upward arrow (↑), downregulated genes with a downward arrow (↓). Log2fold changes are colour coded (colour gradient from highest upregulated in green to downregulated in red). Genes important for endoreduplication are highlighted in blue.

| Gene | Query | Hit/Gene ID | log2FC | up/down |
|---|---|---|---|---|
| CCS52A1 | AT4G22910 | TRINITY_DN89594_c1_g1_i2 | NA | ↑ |
| CCS52A2 | AT4G11920 | | | |
| CCS52B | AT5G13840 | TRINITY_DN67752_c0_g2_i1 | 7.66 | ↑ |
| WEE1 | AT1G02970 | TRINITY_DN103154_c0_g1_i1 | 2.18 | ↑ |
| CDKA;1 | AT3G48750 | TRINITY_DN97304_c1_g1_i1 | NA | ↑ |
| CDKB1;1 | AT3G54180 | TRINITY_DN83815_c0_g1_i1 | 10.71 | ↑ |
| CDKB1;2 | AT2G38620 | | | |
| CDKB2;1 | AT1G76540 | TRINITY_DN100092_c5_g6_i1 | 3.59 | ↑ |
| CDKB2;2 | AT1G20930 | TRINITY_DN100092_c0_g1_i1 | 6.19 | ↑ |
| CDKC;1 | AT5G10270 | TRINITY_DN97810_c3_g2_i1 | NA | ↓ |
| CDKD;1 | AT1G73690 | | | |
| CDKD;3 | AT1G18040 | TRINITY_DN98599_c2_g1_i2 | 3.20 | ↑ |
| CDKF;1 | AT4G28980 | TRINITY_DN95900_c0_g1_i1 | NA | ↑ |
| CKS1 | AT2G27960 | | | |
| CKS2 | AT2G27970 | TRINITY_DN108344_c0_g1_i1 | 3.37 | ↑ |
| CYCA1;1 | AT1G44110 | TRINITY_DN97779_c1_g1_i4 | 2.26 | ↑ |
| CYCA1;2 | AT1G77390 | TRINITY_DN79679_c0_g2_i1 | NA | ↑ |
| CYCA2;1 | AT5G25380 | | | |
| CYCA2;2 | AT5G11300 | TRINITY_DN50245_c0_g3_i1 | 2.37 | ↑ |
| CYCA2;3 | AT1G15570 | TRINITY_DN99644_c2_g2_i2 | 7.95 | ↑ |
| CYCA2;4 | AT1G80370 | TRINITY_DN99644_c2_g3_i8 | 1.85 | ↑ |
| CYCB1;1 | AT4G37490 | TRINITY_DN98655_c3_g8_i1 | 2.35 | ↑ |
| CYCB1;2 | AT5G06150 | TRINITY_DN99804_c2_g1_i1 | 9.33 | ↑ |
| CYCB1;3 | AT3G11520 | TRINITY_DN99804_c3_g1_i1 | 4.07 | ↑ |
| CYCB1;4 | AT2G26760 | | | |
| CYCB2;1 | AT2G17620 | TRINITY_DN35954_c0_g2_i1 | NA | ↑ |
| CYCB2;2 | AT4G35620 | TRINITY_DN92473_c1_g2_i1 | NA | ↑ |
| CYCB2;3 | AT1G20610 | TRINITY_DN75673_c0_g2_i1 | NA | ↑ |
| CYCB2;4 | AT1G76310 | TRINITY_DN64261_c0_g1_i1 | 3.54 | ↑ |
| CYCB3;1 | AT1G16330 | TRINITY_DN88505_c1_g1_i2 | 8.19 | ↑ |
| CYCD1;1 | AT1G70210 | TRINITY_DN84213_c1_g2_i1 | 2.83 | ↑ |
| CYCD2;1 | AT2G22490 | TRINITY_DN94583_c1_g1_i1 | 1.62 | ↑ |
| CYCD3;1 | AT4G34160 | TRINITY_DN93529_c0_g2_i6 | 4.25 | ↑ |
| CYCD3;2 | AT5G67260 | TRINITY_DN3146_c0_g1_i1 | NA | ↑ |
| CYCD3;3 | AT3G50070 | TRINITY_DN70625_c0_g1_i1 | 1.58 | ↑ |
| CYCD5;1 | AT4G37630 | TRINITY_DN30164_c0_g2_i1 | 1.78 | ↑ |
| CYCD6;1 | AT4G03270 | TRINITY_DN3457_c1_g1_i1 | 2.66 | ↑ |
| DPa | AT5G02470 | TRINITY_DN98378_c3_g1_i2 | NA | ↑ |
| E2Fa | AT2G36010 | TRINITY_DN99974_c0_g1_i8 | NA | ↑ |
| E2Fc | AT1G47870 | TRINITY_DN2252_c1_g1_i1 | NA | ↑ |

| | | | | |
|---|---|---|---|---|
| E2Fd/DEL2 | AT5G14960 | TRINITY_DN63323_c0_g1_i1 | 2.51 | ↑ |
| E2Ff/DEL3 | AT3G01330 | TRINITY_DN83317_c0_g1_i1 | 6.78 | ↑ |
| DEL1 | AT3G48160 | TRINITY_DN91414_c0_g1_i2 | 6.0502 | ↑ |
| KRP1 | AT2G23430 | | | |
| KRP3 | AT5G48820 | TRINITY_DN93265_c1_g2_i1 | NA | ↑ |
| KRP5 | AT3G24810 | TRINITY_DN66009_c0_g1_i1 | NA | ↓ |
| KRP7 | AT1G49620 | | | |
| MYB3R1 | AT4G32730 | TRINITY_DN100421_c0_g4_i1 | 5.17 | ↑ |
| MYB3R4 | AT5G11510 | TRINITY_DN92052_c1_g1_i1 | 1.93 | ↑ |
| SIM | AT5G04470 | TRINITY_DN47290_c1_g1_i1 | NA | ↓ |
| SMR1 | AT3G10525 | TRINITY_DN89587_c0_g1_i1 | NA | ↓ |
| SMR10 | AT2G28870 | TRINITY_DN77455_c0_g2_i1 | -1.55 | ↓ |
| SMR11 | AT2G28330 | | | |
| SMR13 | AT3G20898 | | | |
| SMR14 | AT5G59360 | | | |
| SMR2 | AT1G08180 | TRINITY_DN159569_c1_g1_i1 | -1.59 | ↓ |
| SMR3 | AT5G02420 | | | |
| SMR6 | AT5G40460 | | | |
| SMR8 | AT1G10690 | TRINITY_DN90258_c0_g1_i1 | -1.56 | ↓ |
| SMR9 | AT1G51355 | | | |
| RBR1 | AT3G12280 | TRINITY_DN97437_c1_g3_i2 | 1.39 | ↑ |
| CDC20 | AT4G33270 | TRINITY_DN94967_c0_g1_i3 | 7.61 | ↑ |
| APC1 | AT5G05560 | TRINITY_DN99913_c2_g1_i2 | NA | - |
| APC2 | AT2G04660 | TRINITY_DN99749_c1_g2_i4 | NA | ↑ |
| APC6 | AT1G78770 | TRINITY_DN69002_c0_g1_i1 | 1.12 | ↑ |

**Supplementary Table 2.** Cell cycle genes (list from Bothwell et al. 2010) differentially expressed in *Maullinia ectocarpii* infected *Ectocarpus siliculosus* (Ec32m) in comparison to noninfected *Ectocarpus siliculosus*. The log two-fold changes (log2FC) are colour coded, with green as the highest value and red the lowest value. The upregulated genes are indicated with an upward arrow (↑) whilst the downregulated genes are highlighted with a downward arrow (↓). Genes thought to be important for the switch from the mitotic cell cycle to the endocycle are highlighted in blue.

| Gene | Query | Hit/Gene ID | log2FC | significant | up/down |
|---|---|---|---|---|---|
| Ectsi FZR1 (CDH1-Ccs52) | esi0012_0096 | Ec-15_000510 | 0.04 | NO | ↑ |
| Ectsi Wee1 | esi0495_0011 | Ec-00_006420 | -1.42 | YES | ↓ |
| Ectsi CDKA1 | esi0037_0007 | Ec-04_005630 | 0.81 | YES | ↑ |
| Ectsi CDKA2 (CDKB) | esi0041_0093 | Ec-02_003750 | -0.26 | NO | ↓ |
| Ectsi CDKB/4-like | esi0129_0022 | Ec-05_004440 | 0.20 | NO | ↑ |
| Ectsi CDKC1 | esi0007_0143 | Ec-08_002250 | 0.49 | YES | ↑ |
| Ectsi CDKC2,1 | esi0073_0098 | Ec-10_001350 | -0.30 | YES | ↓ |
| Ectsi CDKC2,2 | esi0010_0208 | Ec-20_004690 | 0.79 | YES | ↑ |
| Ectsi CDKD1 | esi0236_0041 | Ec-04_000570 | 0.59 | YES | ↑ |
| Ectsi CDKI1 | esi0191_0048 | Ec-13_000360 | -0.22 | NO | ↓ |
| Ectsi CDKH1 | esi0011_0074 | Ec-03_002400 | 0.53 | YES | ↑ |
| Ectsi CDK-related | esi0011_0004 | Ec-03_002760 | 0.34 | NO | ↑ |
| Ectsi CKS1 | esi0085_0076 | Ec-06_000360 | -0.37 | NO | ↓ |
| Ectsi CKS2 | esi0401_0014 | Ec-06_003970 | -2.10 | YES | ↓ |
| Ectsi CYCA1 | esi0228_0024 | Ec-01_000740 | -1.09 | YES | ↓ |
| Ectsi CYCB1 | esi0071_0052 | Ec-11_004920 | -1.86 | YES | ↓ |
| Ectsi CYCB2 | esi0295_0026 | Ec-01_009390 | -1.64 | YES | ↓ |
| Ectsi CYCD1 | esi0148_0011 | Ec-07_005740 | -4.80 | YES | ↓ |
| Ectsi CYCD2 | esi0220_0004 | Ec-11_004300 | 0.36 | NO | ↑ |
| Ectsi CYCD3 | esi0070_0096 | Ec-12_004410 | -0.20 | NO | ↓ |
| Ectsi CYCF1 | esi0057_0093 | Ec-07_000220 | 0.71 | YES | ↑ |
| Ectsi CYC | esi0064_0091 | Ec-06_003040 | -0.07 | NO | ↓ |
| Ectsi CYCH | esi0069_0006 | Ec-12_000030 | 0.47 | NO | ↑ |
| Ectsi CYCL1 | esi0091_0056 | Ec-22_002580 | -0.92 | YES | ↓ |
| Ectsi CYCT1 | esi0037_0063 | Ec-04_005240 | 0.07 | NO | ↑ |
| Ectsi CYCT2,1 | esi0119_0017 | Ec-11_001180 | 0.54 | YES | ↑ |
| Ectsi CYCT2,2 | esi0290_0030 | Ec-09_001090 | NA | NO | |
| Ectsi DP | esi0063_0071 | Ec-14_006710 | -0.06 | NO | ↓ |
| Ectsi E2F | esi0014_0069 | Ec-21_005690 | 0.53 | NO | ↑ |
| Ectsi DEL | esi0250_0028 | Ec-17_001960 | -0.25 | NO | ↓ |
| Ectsi RBR | esi0089_0003 | Ec-14_002750 | -1.18 | YES | ↓ |
| Ectsi CDC20 | esi0047_0070 | Ec-16_002010 | -0.81 | NO | ↓ |
| Ectsi APC1 | esi0182_0021 | Ec-14_001510 | -0.32 | NO | ↓ |
| Ectsi APC2 | esi0044_0151 | Ec-04_003800 | | | |
| Ectsi APC3 | esi0331_0027 | Ec-03_001570 | 0.04 | NO | ↑ |
| Ectsi APC4 | esi0347_0010 | Ec-26_002710 | 0.27 | NO | ↑ |
| Ectsi APC5 | esi0255_0019 | Ec-06_009310 | 0.22 | NO | ↑ |
| Ectsi APC6 | esi0043_0021 | Ec-05_003300 | -0.42 | NO | ↓ |
| Ectsi APC7 | esi0203_0024 | Ec-01_008970 | 0.23 | NO | ↑ |
| Ectsi APC8 | esi0160_0056 | Ec-03_005170 | 0.06 | NO | ↑ |
| Ectsi APC10 | esi0035_0106 | Ec-19_002640 | 0.85 | YES | ↑ |
| Ectsi APC11 | esi0327_0018 | Ec-07_003050 | | | |

**Supplementary Table 3.** *Maullinia ectocarpii* genes related to the cell cycle (cog cat = D) which are predicted as effectors (EffectorP).

| Gene | Gene expression | EffectorP | eggNOG | | | uniprot | | blastp |
|---|---|---|---|---|---|---|---|---|
| ID | TPM | Prediction | COG cat | predicted_gene | annotation | protein | domain | % identity |
| TRINITY_DN12131_c1_g1_i1 | 608.51 | Effector | D | ANAPC10 | complex subunit 10 | Anaphase-promoting complex subunit 10 | DOC | 67.4 |
| TRINITY_DN35950_c0_g1_i1 | 560.22 | Effector | D | DDB1A | Protein phosphatase 6, regulatory subunit | Cysteine dioxygenase | | 31.8 |
| TRINITY_DN32460_c0_g1_i1 | 217.38 | Effector | D | MOB1B | MOB kinase activator | Mps one binder kinase activator-like 1 protein | | 65.6 |
| TRINITY_DN28765_c0_g2_i1 | 83.14 | Effector | D | MOB1 | MOB kinase activator | Uncharacterized protein | | 76.6 |
| TRINITY_DN34562_c0_g1_i1 | 50.15 | Effector | D | NA | MOB kinase activator | uncharacterized Protein | | 57.5 |
| TRINITY_DN36819_c3_g2_i1 | 8.46 | Effector | D | NBP35 | Component of the cytosolic iron-sulfur (Fe S) protein assembly (CIA) machinery. Required for maturation of extramitochondrial Fe-S proteins. The NBP35-CFD1 heterotetramer forms a Fe-S scaffold complex, mediating the de novo assembly of an Fe-S cluster and its transfer to target apoproteins | Cytosolic Fe-S cluster assembly factor NUBP1 homolog | | 66.8 |
| TRINITY_DN28765_c0_g1_i1 | 4.27 | Effector | D | MOB1 | MOB kinase activator | Uncharacterized protein | | 84.0 |
| TRINITY_DN31047_c0_g1_i1 | 2.29 | Effector | D | CHEK2 | serine threonine-protein kinase | Protein kinase domain-containing protein | Protein kinase | 51.9 |
| TRINITY_DN26027_c0_g2_i1 | 1.44 | Effector | D | DCLK3 | doublecortin-like kinase | Protein kinase domain-containing protein | Protein kinase | 72.3 |
| TRINITY_DN28783_c0_g4_i1 | 0.22 | Effector | D | NA | Cyclin K | Uncharacterized protein | Cyclin | 51.6 |

**Supplementary Table 4.** Cell cycle related genes in *Plasmodiophora brassicae* which act as potential effectors (predicted by EffektorP) (on their host *Brassica oleracea*).

| **Gene** | **Gene expression** | **EffectorP** | **eggNOG** | | | **kegg** | | **blastp** |
|---|---|---|---|---|---|---|---|---|
| ID | FPKM_YG | Prediction | COG cat | predicted_gene_name | eggNOG annot | KEGG_identifier | KEGG_description | % identy |
| TRINITY_DN122143_c1_g1_i1 | 71.92 | Effector | D | NA | Mitotic checkpoint protein | BUB3 | cell cycle arrest protein BUB3 | 100 |
| TRINITY_DN34123_c0_g1_i1 | 60.08 | Effector | D | AURKA | serine threonine-protein kinase | NA | NA | 98.266 |
| TRINITY_DN34569_c1_g1_i1 | 37.74 | Effector | D | CDC25B | cell division cycle 25 homolog | MIH1 | M-phase inducer tyrosine phosphatase [EC:3.1.3.48] | 100 |
| TRINITY_DN115607_c1_g1_i1 | 26.16 | Effector | D | BRUCE | Baculoviral IAP repeat containing | BIRC6, BRUCE | baculoviral IAP repeat-containing protein 6 (apollon) [EC:2.3.2.23] | 100 |
| TRINITY_DN54734_c0_g1_i1 | 25.52 | Effector | D | NA | MOB family member 4, phocein | NA | NA | 100 |
| TRINITY_DN41093_c0_g2_i1 | 21.31 | Effector | D | APC7 | anaphase promoting complex subunit 7 | APC7 | anaphase-promoting complex subunit 7 | 99.587 |
| TRINITY_DN80052_c0_g2_i1 | 11.34 | Effector | D | TTK | ttk protein kinase | TTK, MPS1 | serine/threonine-protein kinase TTK/MPS1 [EC:2.7.12.1] | 100 |
| TRINITY_DN120696_c1_g1_i1 | 7.39 | Effector | D | WDR74 | WD repeat domain 74 | NA | NA | 100 |
| TRINITY_DN77936_c1_g1_i1 | 7.15 | Effector | D | NA | cysteine | NA | NA | 100 |
| TRINITY_DN11234_c0_g1_i1 | 6.64 | Effector | D | RPTOR | regulatory associated protein of MTOR | RAPTOR | regulatory associated protein of mTOR | 98.745 |
| TRINITY_DN86908_c0_g2_i2 | 5.37 | Effector | D | SMC1 | structural maintenance of chromosomes protein | SMC1 | structural maintenance of chromosome 1 | 100 |
| TRINITY_DN58085_c0_g2_i1 | 5.34 | Effector | D | ANAPC10 | complex subunit 10 | APC10, DOC1 | anaphase-promoting complex subunit 10 | 100 |
